# A new twist in ABC transporter mediated multidrug resistance – Pdr5 is a drug/proton co-transporter

**DOI:** 10.1101/2021.05.26.445758

**Authors:** Manuel Wagner, Daniel Blum, Stefanie L. Raschka, Lea-Marie Nentwig, Christoph G. W. Gertzen, Minghao Chen, Christos Gatsogiannis, Andrzej Harris, Sander H. J. Smits, Richard Wagner, Lutz Schmitt

## Abstract

The two major efflux pump systems are involved in multidrug resistance (MDR): (i) ATP binding cassette (ABC) transporters and (ii) secondary transporters. While the former use binding and hydrolysis of ATP to facilitate export of cytotoxic compounds, the latter utilize electrochemical gradients to expel their substrates. Pdr5 from *Saccharomyces cerevisiae* is a prominent member of eukaryotic ABC transporters that are involved in MDR and used as a frequently studied model system. Although investigated for decades, the underlying molecular mechanisms of transport and specificity remain elusive. Here, we provide electrophysiological data on reconstituted Pdr5 demonstrating that this MDR efflux pump does not only actively translocate its substrates across the lipid bilayer, but generates a proton motif force in the presence of Mg^2+^-ATP and substrates by acting as a proton/drug co-transporter. Importantly, a strictly substrate dependent co-transport of protons was also observed in *in vitro* transport studies using Pdr5-enriched plasma membranes. Similar observations have not yet been reported for any other MDR efflux pump. We conclude from these results that the mechanism of MDR conferred by Pdr5 and likely other transporters is more complex than the sole extrusion of cytotoxic compounds and involves secondary coupled processes suitable to increase the effectiveness.

## Introduction

The phenomenon of multidrug resistance (MDR) exists in all cells ranging from prokaryotes to higher eukaryotes and describes the ability to prevent the efficacy of a wide range of functionally and structurally different cytotoxic compounds. One of the major mechanisms that confer MDR is the overexpression of certain efflux pumps expelling these toxic compounds into the extracellular space (1). Two classes of transporters are involved: (1) ATP binding cassette (ABC) transporters that utilize the binding and hydrolysis of ATP to transport their substrates and (2) secondary transporters of the major facilitator superfamily (MFS) as well as the RND and MATE family that use an existing electrochemical gradient across the membrane in order to export these compounds (2–5). Although studied for decades, the molecular mechanisms underlying the resistance conferred by these membrane proteins remain elusive in most cases (6).

In general, membrane transporters and channels are characterized by a high level of substrate specificity. Therefore, the large variety of substrates of individual MDR transporters raises the question whether these MDR transporters directly transport such a broad variety of structurally unrelated compounds or whether their physiological role within the cellular membrane indirectly protects the cell against cytotoxic compounds (2, 7). For the well-characterized human MDR ABC transporter P-gp (MDR1 or ABCB1), it was demonstrated that cells expressing this protein have an altered cytosolic pH and membrane potential compared to cells lacking MDR1. Earlier, *in vivo* studies revealed that Cdr1, a major MDR ABC transporter from pathogenic *Candida albicans* and a close homologue of *S. cerevisiae* Pdr5, alters the extracellular pH (8). Similar to MDR1, studies reported alterations of the membrane environment through altered ion gradients by MDR efflux pumps. In a study of the isolated transmembrane domain of the bacterial multidrug transporter LmrA from *Lactococcus lactis*, it was demonstrated that the transporter still facilitates proton motive force (PMF) driven uptake of drugs in absence of the nucleotide binding domains (NBDs) (9). This finding was later confirmed for full-length LmrA, which, however, requires that the physiological PMF results in the uptake of drugs through this membrane protein (10). The complexity of ion translocation by ABC transporters becomes more evident, if two other studies regarding this protein are taken into consideration. The first one demonstrated that LmrA is an ethidium bromide - H^+^ - Na^+^ - Cl^−^ symporter, while more recently it was shown that it can act as a Hepes^+^ - H^+^ - Cl^−^/2Na^+^ antiporter (11, 12). Another study involving the ABC transporter MsbA reported the requirement of the PMF for this membrane protein, as ATP hydrolysis alone was insufficient to drive substrate translocation (13).

In our study, we used purified Pdr5, a long and intensively-studied member of the MDR ABC transporter family and the major player of the pleiotropic drug resistance (PDR) network of *S. cerevisiae* (14–18). Like other MDR ABC efflux pumps, *in vivo* and *in vitro* studies with plasma membrane vesicles clearly established that Pdr5 confers resistance towards a wide variety of hydrophobic compounds that are structurally unrelated, as well as metabolites (15, 17, 19). Only recently it was possible to isolate this transporter and purify it in an active state (20). Based on this possibility, we reconstituted Pdr5 into planar lipid bilayers and provide *in vitro* data demonstrating that Pdr5 actually co-transports drug/protons in a strictly ATP- and substrate-dependent manner. These results were supported by *in* vitro transport studies using plasma membranes highly enriched in Pdr5. Importantly, we show that contrary to LmrA or MsbA, Pdr5 does not rely on the presence of a PMF as an energy source, rather generates itself a PMF across the membrane in the presence of ATP and substrate. We conclude that the substrate/ATP induced PMF by Pdr5 serves as a self-activation representing a new mechanism of MDR.

## Results

### *In vitro* transport assay for Pdr5

In our efforts to reconstitute purified wild-type Pdr5 (Pdr5_WT_) into synthetic liposomes and to establish an *in vitro* transport assay, we noticed robust ATPase activity comparable to the activity observed in detergent micelles (20). In plasma membrane preparations containing Pdr5_WT_, a robust, real-time rhodamine 6G (R6G) transport assay is often used to study Pdr5_WT_ (17, 18). This assay is schematically summarized in Figure 1A. Here, plasma membranes enriched in Pdr5_WT_ likely present as inside-out vesicles, although the precise nature (sheets or vesicles) of the plasma membrane is unknown, are used. Incubation of plasma membrane vesicles enriched in Pdr5_WT_ with R6G results in a nearly quantitative insertion of R6G into the bilayer, followed by a slower equilibration between inner and outer leaflets (indicated as “1” in Figure 1A). Addition of ATP (“2”) to the Mg^2+^ containing buffer results in energization of Pdr5_WT_ and pumping of R6G from the outer to the inner leaflet, which decreases the R6G fluorescence intensity (Δ*F_i_*). Normally, a decrease on the order of 50-80% is achieved, which is due to oligomer formation that are non-fluorescent (17, 21). As soon as the active, Pdr5_WT_ mediated transport and the spontaneous back flip of R6G reach equilibrium, no macroscopic change in the fluorescence intensity is observed (“3”). Addition of, for example, EDTA (“4”), which complexes the cofactor Mg^2+^, restores the R6G fluorescence to its initial value prior to the addition of ATP (“5” in Figure 1A).

**Figure 1.**
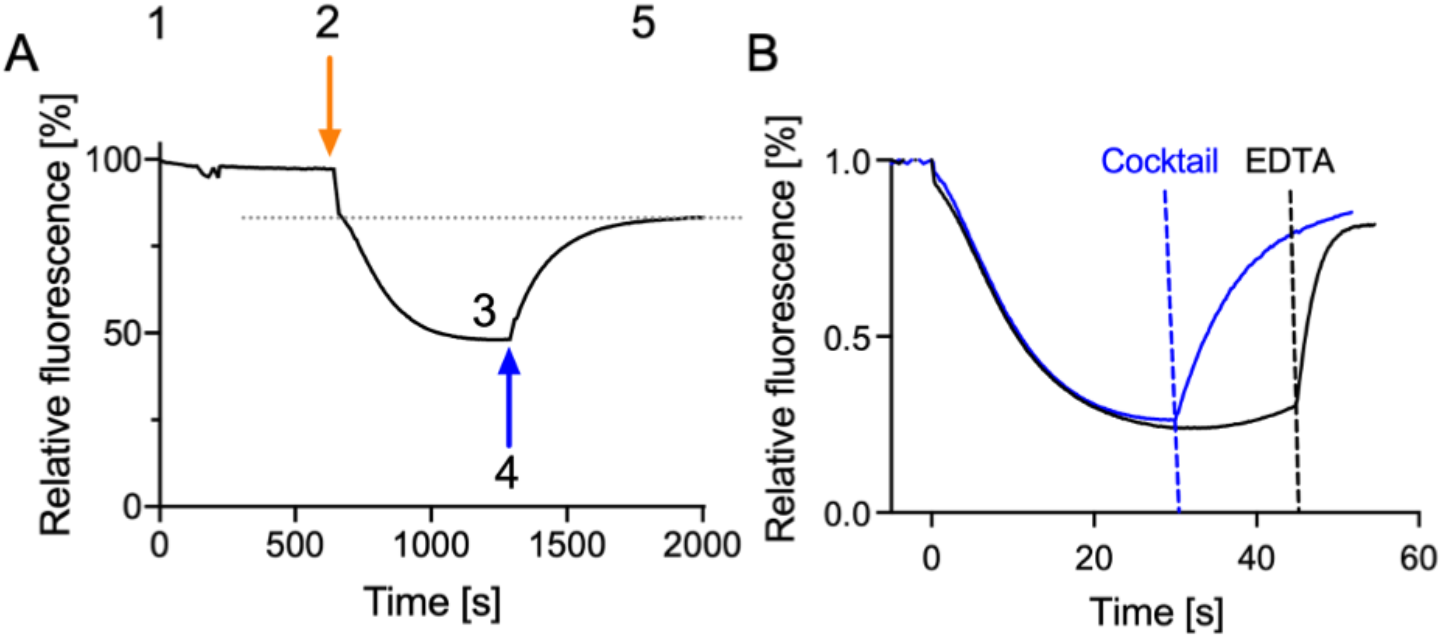
R6G transport assay. (A): Real-time R6G transport in Pdr5WT enriched plasma membranes. At time point 0, R6G is added to a solution containing yeast plasma membranes (“1”). After energization of Pdr5_WT_ by addition of ATP (“2”), R6G is transported from the outer to the inner leaflet until a new equilibrium value of the fluorescence intensity is reached (“3”). The Pdr5_WT_-mediated transport results in the formation of non-fluorescent R6G oligomers in the inner leaflet as indicated by the intensity drop of approximately 60%. At equilibrium, active transport of R6G by Pdr5_WT_ and spontaneous flip-back reach steady state. At “4” the inhibitor EDTA, which complexes the essential cofactor Mg^2+^, is added, which results in restoration of the initial fluorescence intensity (“5”) which is highlight by the gray, dashed line. **(B):** Dependence of R6G fluorescence intensity in plasma membrane preparations highly enriched in Pdr5_WT_ on ATP induced R6G transport in the absence (control) and the presence of an artificial K^+^ gradient. (Control): R6G transport was initiated by addition of 5 mM ATP-Mg^2+^, at the indicated time, EDTA (1 mM) was added. (K^+^ gradient): After addition of 100 mM KCl and 5 mM ATP-Mg^2+^, nigericin (5 μM), valinomycin (0.18 μM) and gramicidin (2 μg/ml) (“Cocktail”) were added at the indicated time.

In contrast, in Pdr5_WT_ containing proteoliposomes such a decrease was rarely observed. Only in less than 10% of the attempts, Δ*F_i_* as read-out for substrate transport was detected with Δ*F_i_* < 10%, not comparable to the more than 50-80% decrease in fluorescence intensity observed when employing plasma membrane preparations (17, 18). In light of the data available for LmrA and MsbA (11–13), we wondered whether ion gradients might also have an impact on Pdr5_WT_-mediated R6G transport. As a first step, we prepared plasma membrane preparations enriched with Pdr5_WT_ (18) and created an artificial potassium and/or proton gradient by either adding 100 mM KCl and valinomycin to the buffer (K^+^ gradient). In the case of the K^+^ gradient, R6G transport was started by addition of 5 mM ATP/Mg^2+^ (Figure 1B). After the transport induced Δ*F_i_* reached a nearly steady state, EDTA was added to stop transport leading to nearly complete reversal of Δ*F_i_*. Next, we tested whether a complete short-circuit of the membrane potential across the plasma membrane during the ATP/Mg^2+^ induced transport might produce a similar effect on Δ*F_i_* as EDTA before. For this, after the R6G transport induced decrease of the fluorescence intensity reached a nearly steady state, nigericin, valincomycin and gramicidin (“Cocktail”) were added in low μM concentrations to suppress the membrane potential completely (Figure 1B) (22, 23). Remarkably, the short circuit of the membrane potential almost completely reversed Δ*F_i_* similar to EDTA (22). These results demonstrated that the fluorescence of R6G is sensitive to electrochemical gradients (PMF) across the membrane. This confers with the known sensitivity of R6G fluorescence towards pH (24) and the membrane potential as derived from *in vivo* and *in vitro* studies (25–27). This also suggests that the decrease in R6G fluorescence in plasma membrane preparation enriched in Pdr5_WT_ might not solely be due to formation of non-fluorescent dimers and trimers (21), but also to changes in the membrane potential.

### Pdr5_WT_ shows voltage dependent ion channel activity

In order to verify whether Pdr5_WT_ activity is modulated by an existing ion gradient or may facilitate voltage activated ionic currents across a membrane, purified Pdr5 was reconstituted into planar lipid bilayer (DOPC/DOPE, 7:3) and voltage dependent currents across the bilayer were monitored. The high purity of the protein used in the electrophysiological experiments was confirmed by MS analysis (see Table S1). In addition to the control experiments with MOK-samples, the same data set as for Pdr5_WT_ was also collected for the ATPase deficient E1036Q mutant (Pdr5EQ) (18).

The slope conductance of the control (“empty”) bilayer was G_sl_ = 3.2 pS (Figure 2A) when a voltage ramp (V_h_ = −100 mV to +100 mV, see Figure 2F top) was applied under symmetrical cis/trans conditions (250 mM KCl, 10 mM HEPES pH 7.0). However, after addition of 2 μl detergent solubilized Pdr5_WT_ (0.6 mg/ml) to the cis-compartment with brief (2min) gentle stirring the voltage induced current across the bilayer increased significantly and a slightly rectifying current voltage relation was observed (Figure 2B). Addition of MOK samples did not show any effect on the i/v-curve. Moreover, repeated aliquot addition (10 times) of the Pdr5 protein to the cis compartment resulted in a large increase (~10^3^-times) in the membrane currents as compared to the control (Figure 2C). Application of voltage gates on the same bilayer as in Figure 2B also showed a significant increase of the membrane currents as compared to the control (Figure 2D) and the resulting plot of the average current (*ī*) versus voltage (Figure 2E) resembled closely the i/v-ramp in Figure 2B. These results give clear evidence that reconstituted Pdr5 facilitates transport of *K*^+^ and *Cl*^−^ ions at comparable high rates, which are also dependent on the amount of reconstituted Pdr5_WT_. Similar results have been obtained for the ATPase deficient E1036Q mutant (Pdr5_EQ_) (see below for details). In order to check whether the high transport rates are caused either by a high number of copies of active Pdr5_WT_ in the membrane or by an “ion channel” mode, we have tested various voltage protocols, which potentially could lead to ion-channel gating of Pdr5_W_.

**Figure 2.**
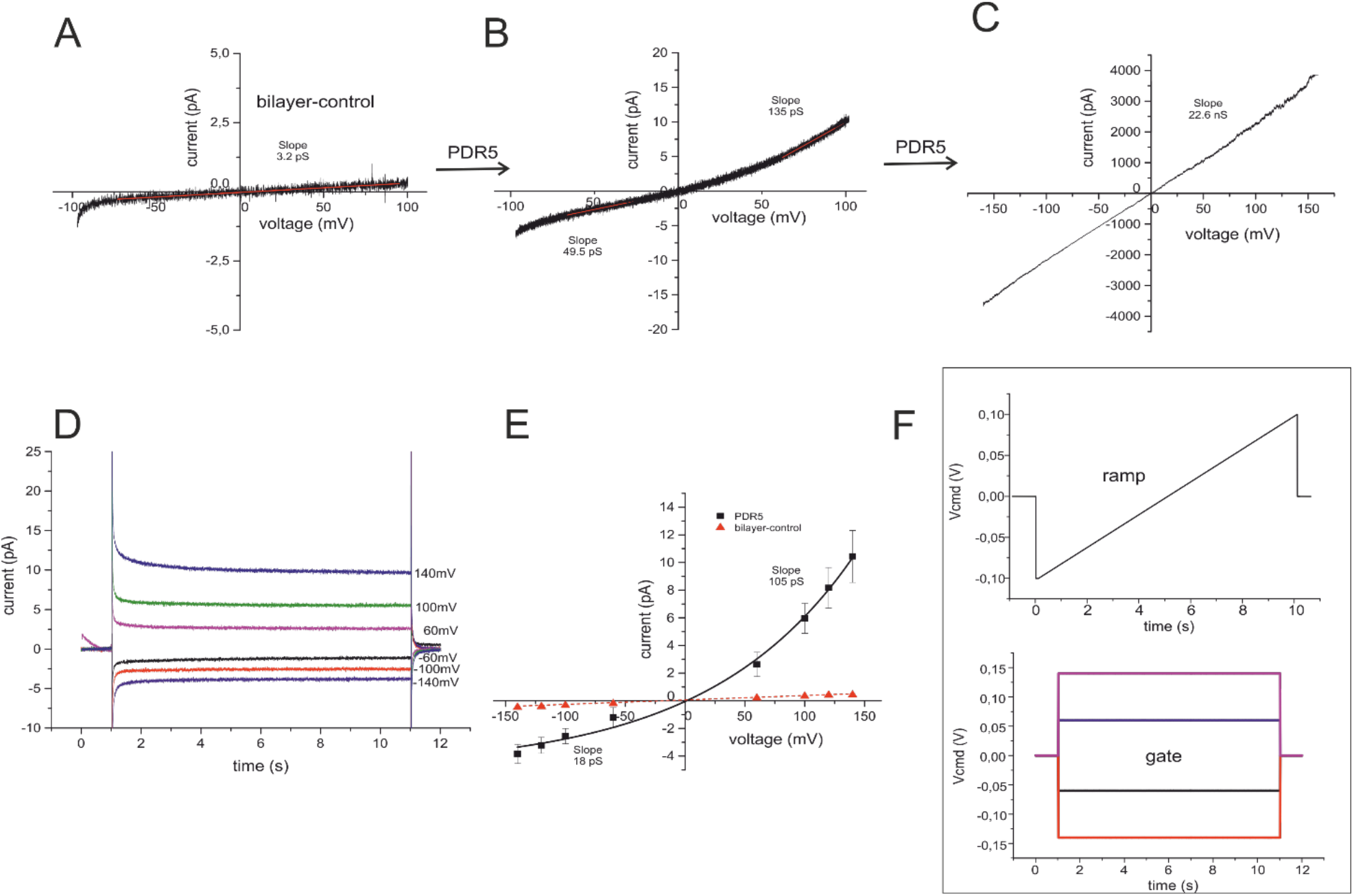
Current recordings from a planar lipid bilayer containing reconstituted Pdr5_WT_. (**A**) Current-voltage ramp measurement with a DOPE/DOPC (7:3) bilayer in symmetrical cis/trans buffer conditions (250 mM KCl, 10 mM HEPES, pH 7.0) (control). (**B**) Current recording of the same bilayer as in (**A**), but after insertion of detergent purified Pdr5_WT_ from the cis compartment using the voltage ramp protocol shown in **(F)** (**C**) Current recording from a bilayer after multiple insertion of detergent purified Pdr5_WT_. (**D**) Current-voltage gates of the same bilayer as in **(B)** using the voltage gate protocol shown in F (lower part). **(E)** Current-voltage relation obtained from the mean currents in **(D)**. (**F**) Applied voltage protocol for voltage-ramps (upper) and voltage gates(lower).

We could indeed find such a voltage-protocol (see Figure 3G, H). Starting from a holding potential of V_h_ = +150 mV for 2-10 min. and using either voltage ramps (Figure 3G) or voltage gates (Figure 3H) we observed a significant increase of the Pdr5_WT_-mediated membrane currents accompanied by brief current flickering (Figure 3B) (G_sl_ = 420pS,). Application of voltage gates allowed for a more detailed analysis of the Pdr5_WT_ mediated current gating (Figure 3C, D, E, F) and Figure S1). The graphs in Figure 3 first clearly show that the reconstituted Pdr5_WT_ mediates fast, flickering current gating with different open channel amplitudes.

**Figure 3.**
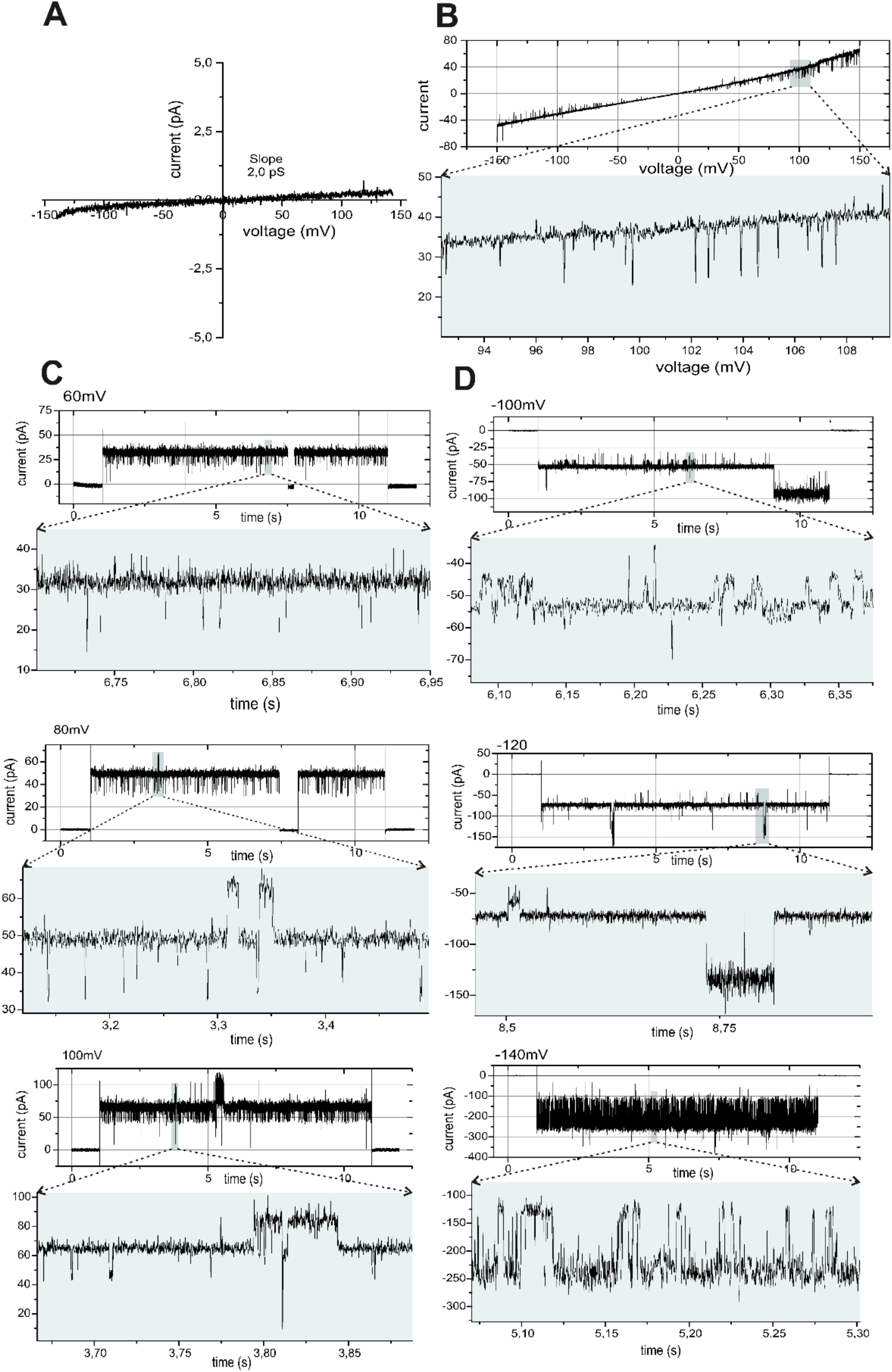
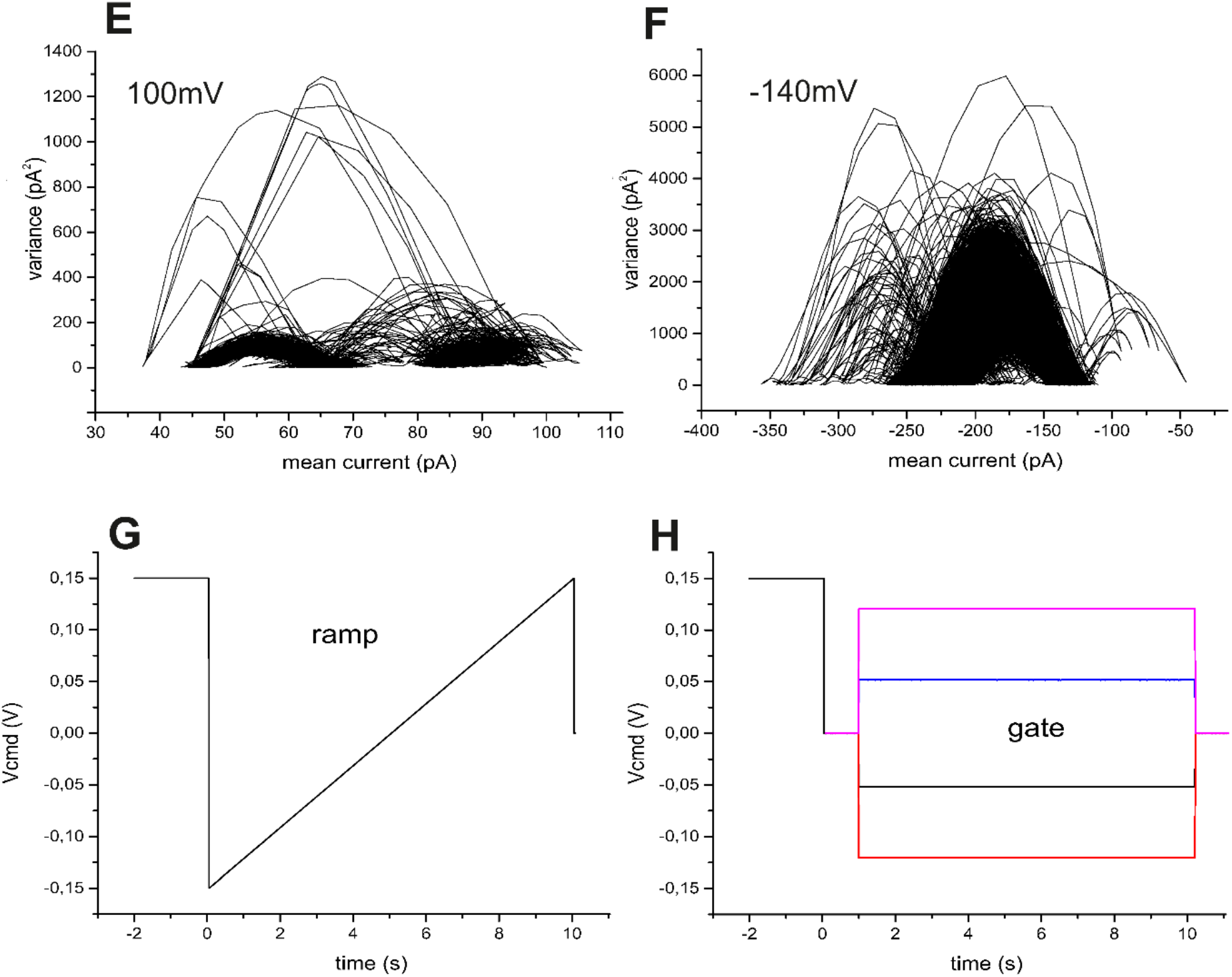
Current recordings with extension plots from a bilayer containing the reconstituted of Pdr5_WT_. **(A)** Current-voltage ramp measurement with a DOPE/DOPC (7:3) bilayer in symmetrical cis/trans buffer conditions (control). (**B**) Current recording of the same bilayer as in (**A**), but after insertion of detergent purified Pdr5_WT_ from the cis compartment using the voltage ramp protocol shown in **(G). (C), (D)** Extension plots of current-voltage gates at the indicated Vh from the same bilayer as in **(B)** using the voltage gate protocol given in **(H) (E), (F)** Mean variance plots for the respective channel recordings shown in **(C)** and **(D)** depicting multiple open channel amplitudes.

Additionally, as indicated by the reproducible rectifying current-voltage relation, Pdr5 incorporation into the membrane occurred almost exclusively unidirectional with the nucleotide binding sites at the cis compartment (for more details see below).

### Pdr5_WT_ possesses three main conductance states

The analysis of the representative current recordings shown in Figure 3, with voltage gates from V_h_ = ±60 mV to V_h_ = ±140 mV show that the observed complex current gating pattern originates from one active channel with up to three open channel amplitudes and not from simultaneous gating of multiple channels (see mean variance analysis in Figure 3E, F and Figure S1). This conclusion is proven by looking closer onto the combined current histogram and mean variance analysis of the current recordings in Figure 3D at V_cmd_ = −140 mV indicating that channel gating occurred with at least 3 different main open channel amplitudes. From the analysis of current recordings at different membrane potentials (V_h_ =±60 mV to V_h_ =±140 mV, see Figure 2C) we collected from 5 different bilayer in a total of 40 recordings all point amplitude histograms (Figure S1). The obtained currents at the given voltages were transformed into conductance values binned in a histogram and the appearing peaks plotted in a box-chart plot (Figure S1C). The following main mean conductance states of Pdr5_WT_ were obtained: G_1_ ≅ 55 ± 24 pS, G_2_ ≅ 304 ± 106 pS, and G_3_ ≅ 665 ± 127 pS. Closer inspection of the time-course of gating events (see Figure 2C and Figure 2D) clearly indicate that one single Pdr5_WT_ unit displayed at least three distinct main subconductance states (for more details see Methods in SI). The selectivity of the Pdr5_WT_ mediated ion currents was tested using asymmetric cis/trans buffer conditions (see Figure S2) and gave a value of *P*_*K*^+^_: *P*_*Cl*^−^_ = 2.6:1.

Remarkably, the mean open state G_2_ was mostly occupied at membrane potentials above *V_cmd_* > ±100 mV. At an ionic strength of 250 mM and using the simplified model of a cylindrical restriction channel region (28, 29) with a length of 1 nm, we calculate from this conductance state the pore diameter of Pdr5 to be d_restriction_ ≅ 6.4 Å.

### ATPase deficient E1036Q mutant (Pdr5_EQ_) reveals similar properties as Pdr5_WT_

After addition of 1 μl detergent solubilized Pdr5_EQ_ (2.0mg/ml) to the cis-compartment with brief (2min) gentle stirring again voltage (voltage ramp starting at V_h_ = 0 mV, see Figure 2F top) induced currents across the bilayer were observed. The amplitude of the open channel currents appeared somewhat smaller as compared to the Pdr5_WT_ with slope conductance of *G_sl_* = 150 pS (-V_cmd_) and *G_sl_* =280 pS (+V_cmd_) were observed (Figure 4A). Again, addition of MOK-samples did not show any effect on the control i/v-curve. Repeated addition (3-times) of the Pdr5 protein to the cis compartment resulted in a large increase (~5-times) in the membrane currents as compared to the control (Figure 4D). Application of voltage gates on the same bilayer as in Figure 4D also showed a significant increase of the membrane currents and the resulting plot of the average current (*ī*) versus voltage (Figure 4E) resembled closely the i/v-ramp in Figure 4D. These results show that that reconstituted Pdr5_EQ_ facilitates transport of *K*^+^ and *Cl*^−^ ions at high rates which are also dependent on the amount of reconstituted Pdr5_EQ_.

**Figure 4.**
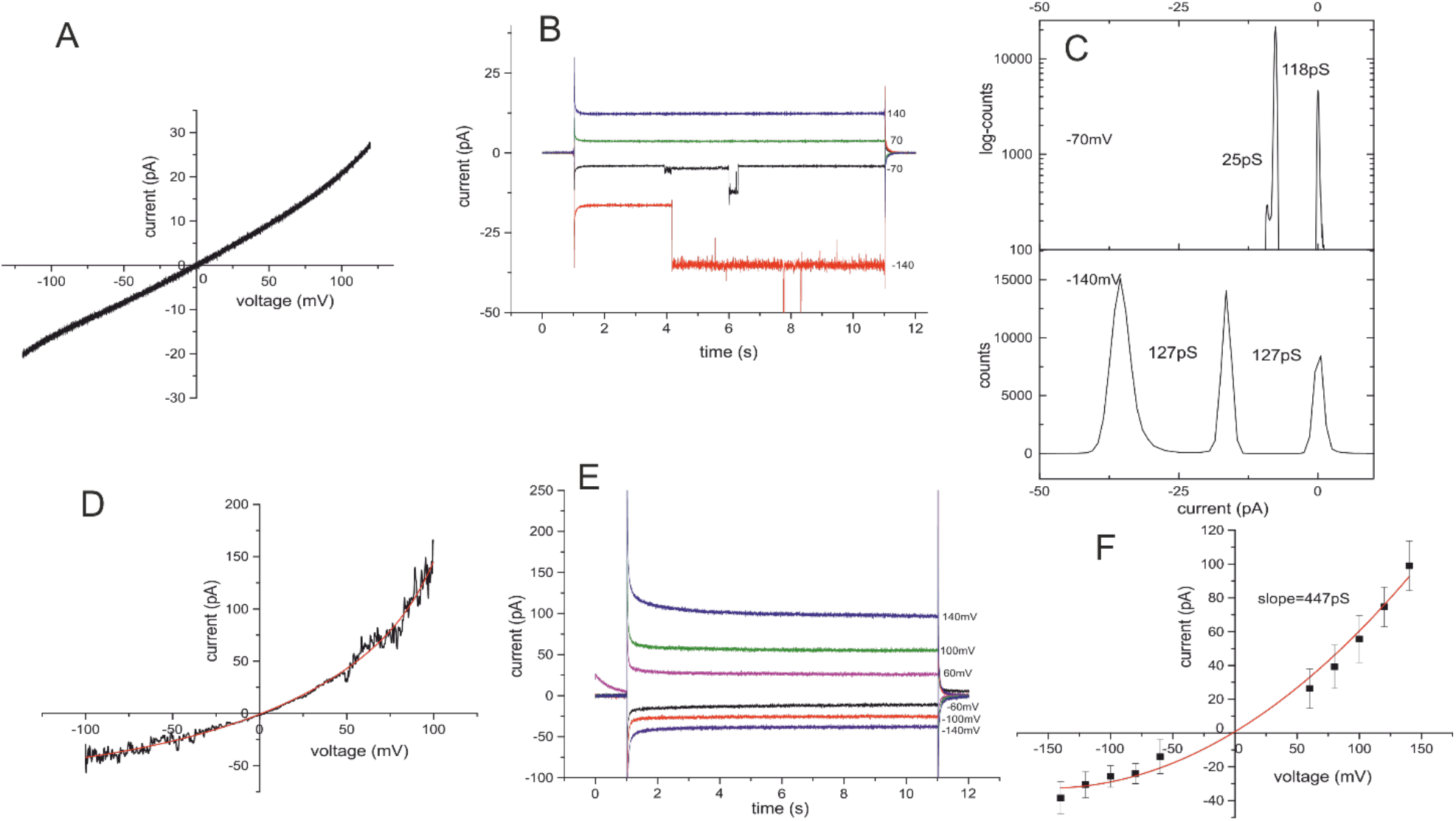
Current recordings from a bilayer containing the reconstituted of Pdr5_EQ_. (**A**) Current-voltage ramp recording with a DOPE/DOPC (7:3) bilayer in symmetrical cis/trans buffer conditions after reconstitution of Pdr5_EQ_ using the voltage ramp protocol shown in Figure 2F. (**B**) Current-voltage gates of the same bilayer as in **(B)**using the voltage gate protocol shown in F (lower part). **(C)** All point amplitude histograms for the recordings shown in **(B)**. **(D)** Current-voltage ramp recording with a DOPE/DOPC (7:3) bilayer in symmetrical cis/trans buffer conditions after multiple additions of Pdr5_EQ_ to the cis compartment using the voltage ramp protocol shown in Figure 2F. (**E**) Current-voltage gates of the same bilayer as in **(D)** using the voltage gate protocol shown in Figure 2F (lower part). **(F)** Current-voltage relation obtained from the mean currents in **(E)**.

However, when the pre-activation command voltage protocol shown in Figure 3G,H was applied to the bilayer containing the reconstituted Pdr5_EQ_ rare current gating events were only observed at negative Vcmd (Figure 4B). From a brief analysis of the current gating events at V_cmd_ = −70 mV to −140 mV, the following main mean conductance states of Pdr5_EQ_ were obtained: G_1_ ≅ 35 ± 12 pS, G_2_ ≅ 120 ± 29 pS. Thus, the current gating properties of the Pdr5_WT_ and the Pdr5_EQ_ mutant are different while voltage activation of Pdr5_WT_ leads to a strong current flickering mainly from the open to the closed state the Pdr5_EQ_ mutant remains mainly in the open state with only low frequent channel closures.

The ion selectivity of the Pdr5_EQ_ mutant as determined in asymmetric cis/trans buffer of *P*_*K*^+^_: *P*_*Cl*^−^_ = 3.5:1 was close to the one of the Pdr5_WT_ (see Figure S2).

### Modulation of Pdr5-mediated ion currents by its substrates

We further tested the effect of known substrates namely R6G, ketoconazole (KA), clotrimazole (CLO) and cycloheximide (CHX) on the voltage-induced ion channel activity of Pdr5_WT_.

Figure 5B shows a current-voltage ramp using the command-voltage protocol shown in the inset of Figure 5A (control, empty bilayer) from a bilayer containing the reconstituted Pdr5_WT_ in symmetrical (cis/trans) buffer. As obvious, ion-conducting activity with frequent channel gating was observed (see enlarged lower panel Figure 5B). The linear slope conductance indicated by the red line was G_sl_= 525 ± 2.4 pS. This value is very close to the Pdr5_WT_ conductance state G3 deduced from channel gating (see above). Therefore, the current-voltage ramp can likely be attributed to a single activated Pdr5_WT_ unit. The brief flickering events resemble the respective channel conductance G1 and G2 in their amplitudes. After addition of 2 mM Mg^2+^-ATP (cis/trans) and 330 nM R6G (1.2 μl 330 μM into 1.2 ml buffer) to the cis compartment of the same bilayer no further voltage dependent channel gating could be observed (Figure 5C). The current was drastically reduced while the form of the current-voltage ramp changed to a more rectifying shape (Figure 5C). In addition, the zero-current crossing (V_rev_), changed completely and unexpectedly from V_rev_ = 0 mV to V_rev_ = −46 mV. This effect was reproducibly observed with an average V_rev_ = −43.8 ± 4.8 mV (n=7) no matter whether the bilayer contained single activated or multiple activated copies of Pdr5_WT_. In this context, it is important to mention that this R6G-induced shift of V_rev_ was strictly dependent on the presence of Mg^2+^-ATP in the cis compartment.

**Figure 5.**
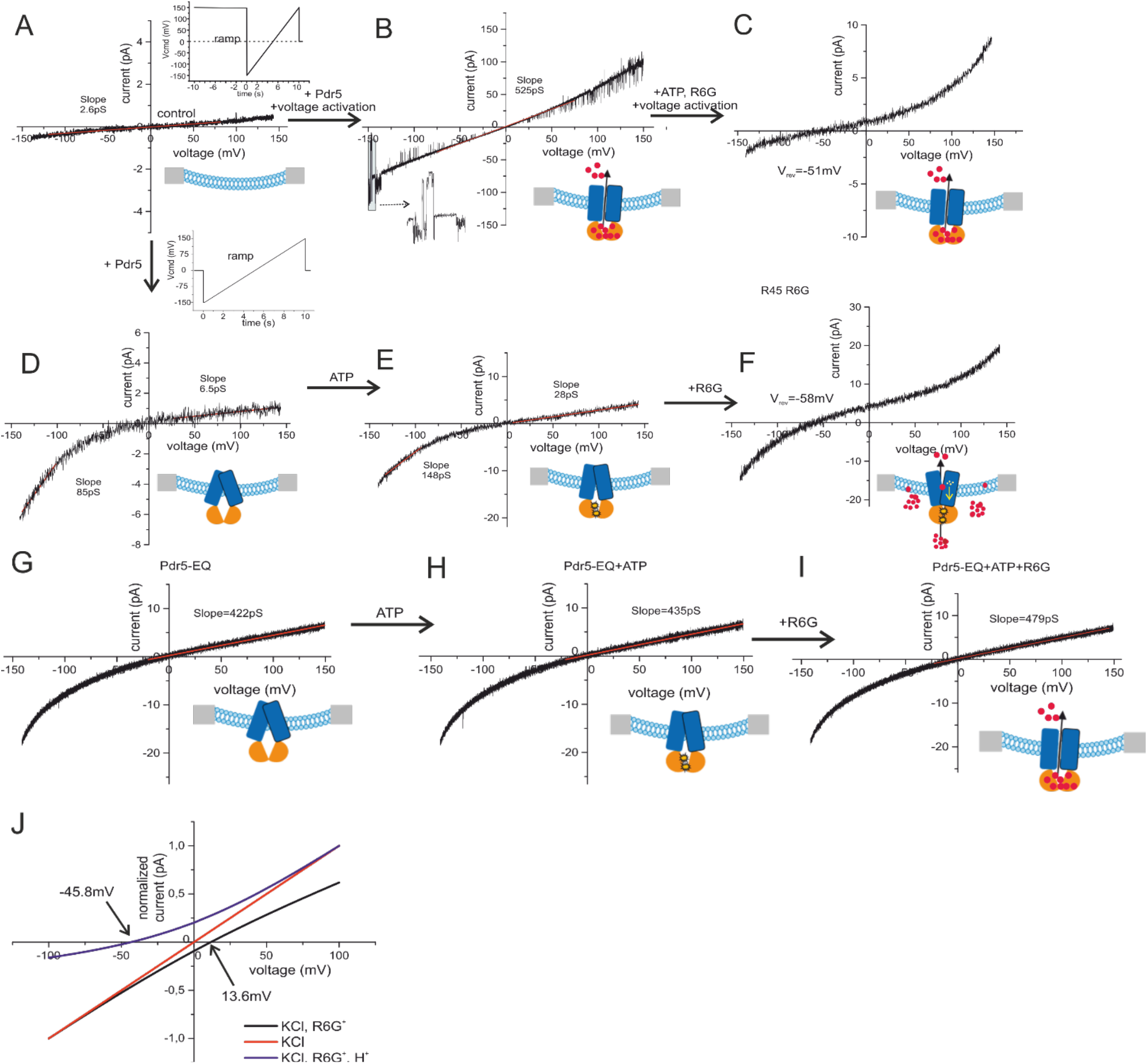
Effects of R6G on Pdr5_WT_ and Pdr5_EQ_ -mediated ion conductance. (**A**) Current-voltage ramp from an empty control bilayer under symmetrical buffer (cis/trans 250 mM KCl, 10 mM HEPES, pH 7.0) **(B)** Current-voltage ramp after reconstitution of Pdr5_WT_ in symmetrical buffer using the command voltage protocol shown in the inset of **(A)**. **(C)** Same bilayer as in **(B)** but after addition (cis) of 1mM Mg^2+^-ATP and 5μM R6G. (**D**) Current-voltage ramp in symmetrical buffer after reconstitution of Pdr5_WT_ using the command voltage protocol shown in the inset below **(A)**. Current-voltage ramp from the same bilayer as in **(D)** but after addition of 1mM Mg^2+^-ATP. **(F)** Same bilayer as in **(E)** but after addition of 5μM R6G. **(G)** Current-voltage ramp in symmetrical buffer (250 mM KCl, 10 mM HEPES, pH 7.0) after reconstitution of Pdr5_EQ_ using the command voltage protocol shown in the inset below **(A)**. **(H)** Current-voltage ramp from the same bilayer as in **(G)** but after addition of 1mM Mg^2+^-ATP (cis/trans). **(I)** Same bilayer as in **(H)** but after addition of 5μM R6G (cis). **(J)** Current-voltage relations calculated by the GHK current equations using applied experimental ionic conditions and the respective permeability (P_x_, (*P*_*K*^+^_/*P*_*Cl*^−^_/*P*_*RG*^+^_)) as variable (see supplemental Information for details) for 3 cases. **Red:** KCl (only permeation of K^+^ and Cl^−^ ions), **Black:** KCl-R6G (permeation of K^+^ and Cl^−^ ions and active transport of R6G^+^ ions), **Blue:** KCl-R6G-H^+^ (permeation of K^+^ and Cl^−^ ions and active co-transport of R6G^+^ and H^+^ ions). Currents were normalized to i_max_. For more details see Methods section and SI.

Addition of Mg^2+^-AMP (trans) (n = 3) did not show this effect. Moreover, in this setup of Pdr5_WT_ reconstituted from the cis compartment, addition of the effectors to the trans compartment (n=3) did also not show any effect. One may argue that the observed shift might be generated due to electrode effects after the addition of 1.2μl 330μM R6G into 1.2ml buffer of the cis compartment only. However, this can be ruled out from both theoretical and experimental considerations. We are using agar protected Ag+/AgCl electrodes (*R_typ_* ≅ 10 *k*Ω), which are carefully checked for electrode offset 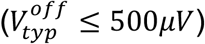, which then is corrected automatically by the amplifier, at the beginning of each bilayer recording (30). In none of the five control experiments with control bilayers, the addition of 1.2 μl 330 μM R6G in EtOH had any effect on the electrode potentials. Moreover, as outlined in more detail below no shift in *V_rev_* was observed in Pdr5_EQ_ containing bilayer at identical experimental conditions. The change in *V_rev_* can therefore only be due to an R6G induced trans-membrane flux of ions as source of the additionally created membrane potential.

In order to explain the observed shift of V_rev_ the macroscopic GHK approach has been widely used. This allows, -based on the ion composition in the cis and trans compartments of the bilayer-, to calculate relative ion fluxes and zero current potentials. Depending on the membrane permeability of the involved ions, the expected reversal potentials for the flux of the given set of ions between the cis and trans compartment separated by a selective permeable membrane can be calculated using the GHK current equation ((31), the details can be found in SI).

The ion concentrations in the cis/trans compartments were kept symmetrical (250 mM KCl, 10 mM HEPES pH 7.0, 2 mM Mg^2+^-ATP) before the addition of 330 nM R6G to the cis compartment and we observed V_rev_ = 0 ± 1.5 mV (n=7). The calculation of the expected V_rev_ by the GHK current equations (see SI for details) before and after addition of charged R6G (charge=+1) clearly showed that the ion concentration at both sides of the Pdr5_WT_ containing membrane can by no means produce a current voltage-relation with a zero-current crossing at V_rev_ = −43.8 ± 4.8 mV (see Figure S3). Therefore, we must conclude that R6G induced a Pdr5_WT_-mediated active ion transport of one of the anions or cations present in solution. With symmetrical 250 mM KCl cis/trans, either Cl^−^ ions would have to be transported from trans to cis to final concentrations of 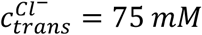 *and* 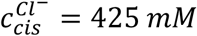, or in the case of K^+^ the ions would be transported from cis to trans to final concentrations of 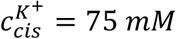 and 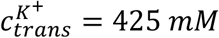 (for details see SI). This massive ion pumping of one of the two ion species would in both cases reduce (Cl^−^) or increase (K^+^) the bulk conductance value in either bilayer compartments, which in fact was not observed. Consequently, transient flux of H^+^ ions is the remaining candidate to build up a membrane potential across the Pdr5_WT_ containing bilayer yielding a zero-current potential of V_rev_ = −43.8 ± 4.8 mV. Introducing H^+^ as an additional ion in the macroscopic GHK current equations we can first determine the direction of the H^+^ flow in our experimental setup as well as the hypothetical macroscopically related changes in the H^+^ concentrations in the cis and trans compartments. The GHK-calculations (details see SI) show that R6G transport is accompanied by a H^+^-transport from cis to trans.

Starting at pH 7 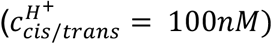 symmetrically, the calculated hypothetical macroscopic changes of the H^+^ concentrations in the cis and trans compartment yield 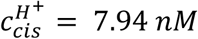 and 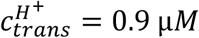 (see Figure 5J and Figure S3 for calculation details). However, since we detect ion flux induced membrane potentials generated by a small number (n<< 100) of channels across a distance of ~20 nm (see SI for details), which requires only tiny amounts of transported charges to establish the membrane potential (see example calculation in SI for details), macroscopic changes of the H^+^ concentrations would not be directly resolvable in our experimental setup.

As is obvious from the calculated current voltage relation with 2mM Mg^2+^-ATP in cis/trans (Figure 5J), the observed shift of *V_rev_* by R6G addition to the cis compartment in Pdr5_WT_ containing bilayer could be explained by active H^+^ transport (H^+^ permeability *P*_*H*^+^_ → ∞) from the cis to the trans compartment. Calculation details for Figure 5J are given in SI.

In similar experiments, we tested whether the non-charged KA, CHX and CLO, other well-known substrates of Pdr5_WT_, also modulate the activity of reconstituted Pdr5_WT_ in planar bilayer. In a series of four different experiments, we observed a Mg^2+^-ATP/KA induced shift of the zero-current potential of V_rev_ = −64 ± 8 mV (n = 4) (see Figure 6).

**Figure 6.**
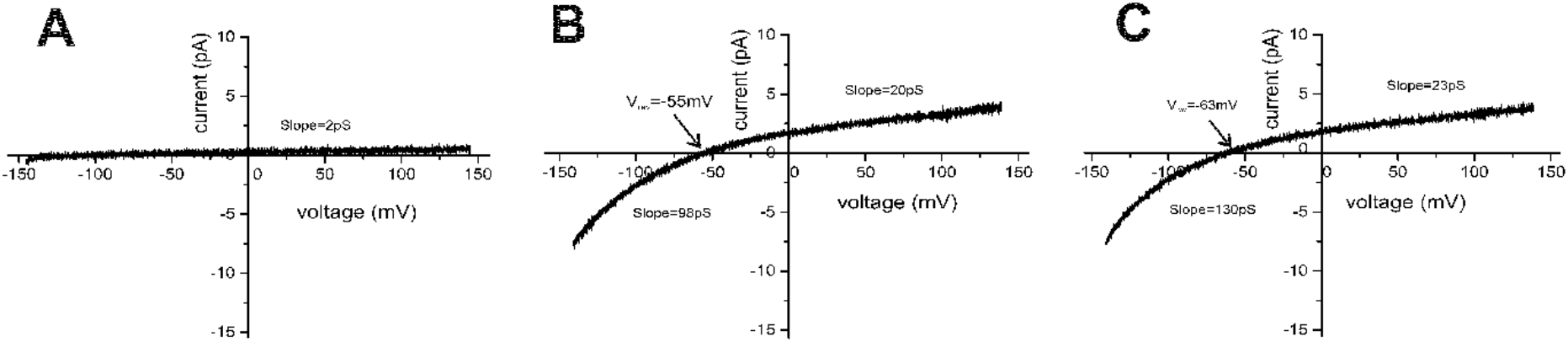
Effects of CLO and KA on Pdr5-mediated ion conductance. (**A**) Control bilayer after reconstitution of Pdr5. (**B**) Same bilayer as in (**A**) but after addition of 1μM CLO to the cis compartment. (**C**) Current-voltage recordings after addition of 330 nM KA to the cis compartment of a Pdr5_WT_ containing bilayer.

Based on these results, we conclude analogously that also KA in the presence of Mg^2+^-ATP induces active Pdr5_WT_-mediated H^+^ transport from the cis to the trans compartment. Interestingly, with CHX as a potential substrate, we did not observe any significant modulation of Pdr5_WT_-mediated currents, in contrast to R6G and KA. However, CLO (1 μM), when added to the cis compartment of a bilayer containing Pdr5_WT_, induced a drastic shift of the reversal potential of V_rev_ = −57 ± 6.7 mV (n=4) in the presence of Mg^2+^-ATP (cis) (see Figure 6).

### The proton gradient is ATPase dependent

In order to verify the significance of the observed modulation of Pdr5_WT_ activity by its substrates, we performed the same set of experiments as describe above except that we used the ATPase deficient Pdr5_EQ_ mutant instead of Pdr5_WT_. This therefore is a proof whether the catalytic ATPase cycle is necessary for the observed shift of V_rev_ and, most importantly, whether the observed ion currents are Pdr5-mediated. As shown before in Figure 4 Pdr5_EQ_ also incorporates into the planar bilayer membrane and mediates voltage dependent ion currents across the membrane. However, Pdr5_EQ_-mediated currents were observed already at membrane potentials ≥ 20 mV. In more than 90% of the attempts during voltage ramps rather large currents were observed (Figure 4A,D) and correspondingly also in voltage gates (Figure 4E, F). These results indicate that a high number of ion conducting Pdr5_EQ_ have been incorporated into the bilayer.

In addition, Pdr5_EQ_ channel gating was rarely observed and the channels remained almost exclusively in the open states as the determined conductance values of the Pdr5_EQ_ were in a similar range as for Pdr5_WT_. This also holds true for the ion selectivity of the Pdr5_EQ_ mutant determined to be *P*_*K*^+^_: *P*_*Cl*^−^_ = =3.5:1 in comparison to 2.6:1 for Pdr5_WT_ (see Figure S2 and SI for details). We also tested whether channel currents of Pdr5_EQ_ could be modulated by the substrates R6G and KA in a similar set of experiments as described above for Pdr5_WT_. We observed that in none out of the 15 attempts with R6G (see Figure 5G-I) or KA any modulation of Pdr5_EQ_-mediated currents showing that ATPase activity and transport of the substrate is an absolute requirement for substrate induced change in *V_rev_*.

### Beauvericin locks Pdr5 in an open state

Beauvericin (BEA) is known as a potent inhibitor of Pdr5’s ATPase and transport activity (32) and additionally can mediate ion transport across the membrane most likely in a carrier like mode (33). We therefore first investigated membrane currents after addition of 0 μM, 3.3 μM and 9.9 μM BEA on the cis side (Figure 7A-C). The conductance values of these measurements are listed in Table S3. Next, we addressed the question whether BEA might suppress the substrate induced H^+^ co-transport of Pdr5_WT_. Figure 7 D-G shows a representative series of experiments with an identical bilayer. The graphs in Figure 7E demonstrates that Pdr5_WT_ was incorporated in the membrane and after addition of R6G to the cis compartment a shift of V_rev_ = − 51 mV was observed. Remarkably, after addition of 1.65 μM BEA the overall current increased significantly and the shift of the reversal potential disappeared completely (Figure 7G).

**Figure 7.**
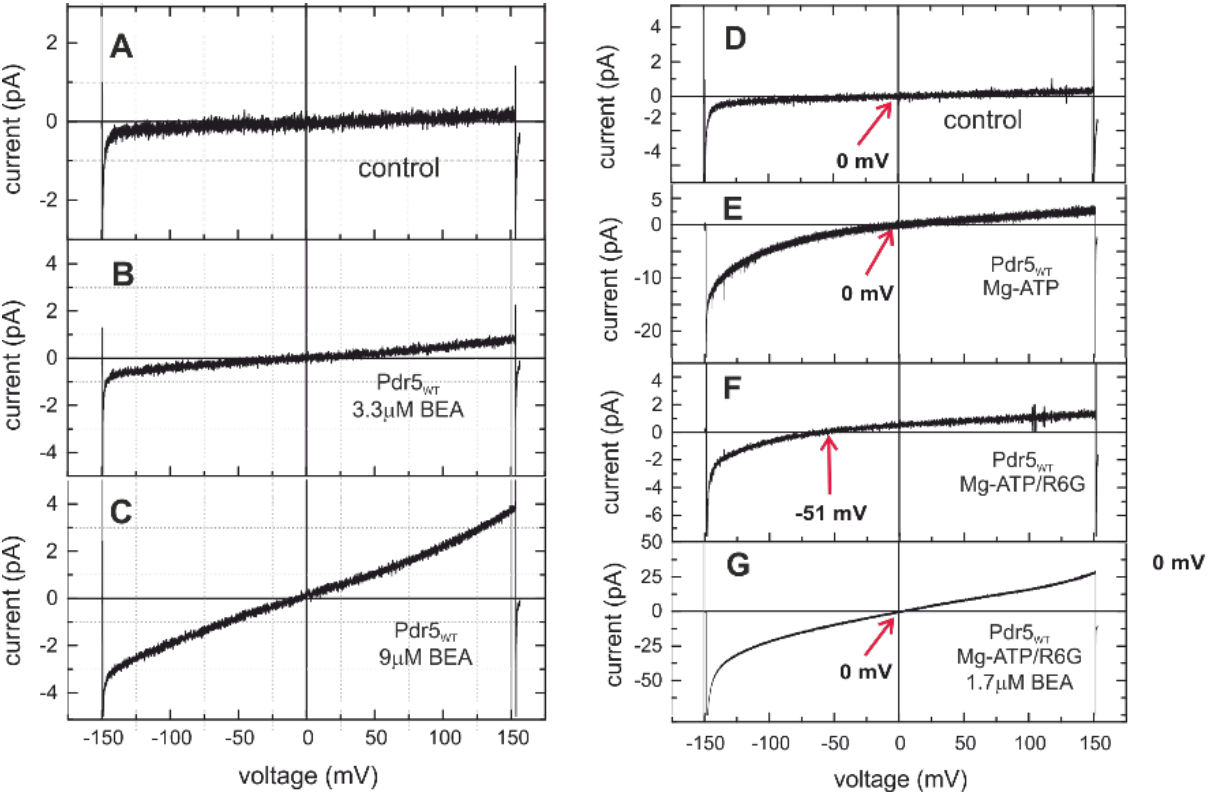
Current-Voltage recording in the presence of BEA in the cis compartment. (**A-C**) Current-voltage ramps with an empty bilayer (**A**) in the presence of 3.3 μM BEA, (**B**), 9.9 μM BEA and (**C**) 0 μM BEA, respectively. (**D-G**) series of current voltage ramps from a single bilayer: (**D**): empty bilayer, (**E**) Pdr5_WT_ incorporated in the presence of Mg^2+^-ATP, (**F**) in the presence of Mg^2+^-ATP/R6G and (**D**) is (**F**) after addition of 3.3 μM BEA.

The conductance values of the different measurements in Figure 7 D-G are listed in Table S4. These recordings show clearly that BEA can act as a carrier, but most importantly acts as a Pdr5 inhibitor that presumably locks Pdr5 in an open channel conformation.

### Molecular docking suggests Beauvericin binding to the Pdr5 substrate binding site

Only recently, structural information of Pdr5 determined by single particle cryo EM became available (34). Thus, we used molecular docking to identify possible binding sites and states of BEA. The above described results suggest that BEA could bind to the R6G binding site to lock Pdr5 in an open state. However, as the starting conformation and placement of the ligand can have an influence on the results of the docking (35), although a small one, we used two different starting placements for BEA.

First, we placed BEA in the binding site of R6G (Figure 8A) as it is the most likely based on our experiments. Second, we placed BEA close to the transmembrane domains (TMDs) in the space between the TMDs and the nucleotide-binding domains (Figure 8B). The rational here was that due to its size BEA might bind to both TMDs while allowing ion fluxes through its macrocycle as an alternative binding mode hypothesis. To include receptor plasticity, we employed four different cryo-EM structures of Pdr5 (34), which show different binding site conformations, after the removal of ligands present in the respective structures. Scoring grids were calculated for at least 40 Å around each initial BEA placements for each of the four Pdr5 conformations and subsequently both initial BEA conformations were docked into each grid. This resulted in 16 valid suggested binding poses. Out of these 16 binding poses nine bind to the R6G binding site in nearly identical conformations (Figure 8C) regardless of their initial placement and center of docking grid. Here, BEA utilizes one of its benzyl moieties such that it binds to a similar location as one of the benzene rings of R6Gs xanthene-like moiety. The five binding poses not binding to the R6G binding site of Pdr5 were either generated in the ADP-orthovanadate bound structure of Pdr5 (four poses) or R6G bound structure (one pose). The ADP-orthovanadate bound structure of Pdr5 shows a shift of one of the transmembrane helices at the R6G binding site. This shift renders the R6G binding site too narrow to accommodate BEA (Figure 8D). As a result, 90 % of all docked BEA binding poses in Pdr5 structures, which could accommodate BEA in the R6G binding site, bind to exactly this binding site in nearly identical conformations. This suggests that BEA binds to the R6G binding site in Pdr5.

**Figure 8.**
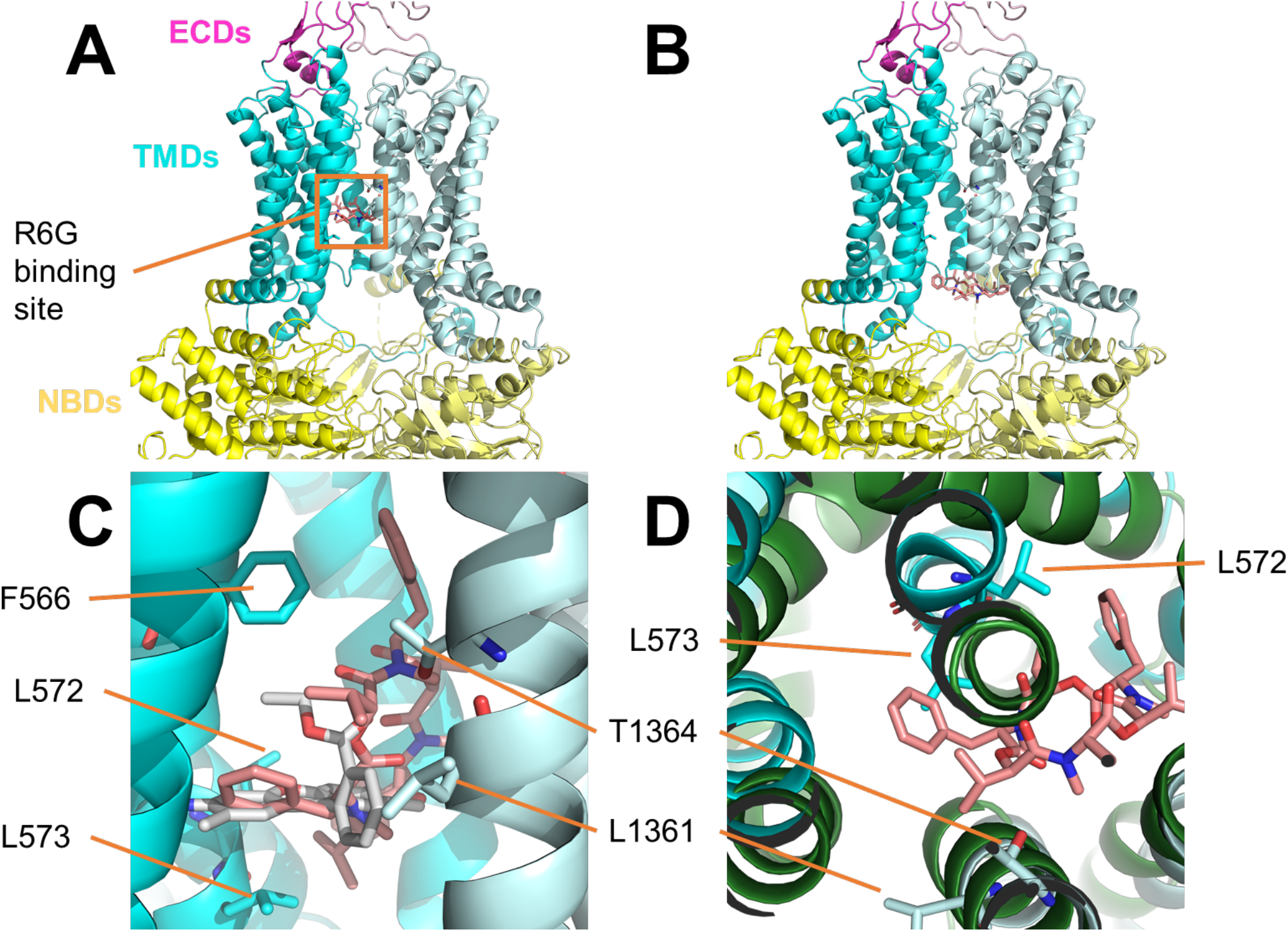
Suggested binding mode of beauvericin (BEA) in Pdr5 identified by molecular docking. (**A, B**) Initial placements and centers for the docking grid generation of BEA in the Rhodamin 6G (R6G) binding site (**A**) and between the TMDs and NBDs (**B**). BEA is shown in salmon sticks and the ectracellular domains (ECDs, magenta shades), transmembrane domains (TMDs, blue shades), and nucleotide binding domains (NBDs, yellow shades) of Pdr5 are shown as cartoon. The R6G binding site is indicated by an orange rectangle. Here the R6G bound structure of Pdr5 was chosen although R6G had been removed for docking and is thus not shown here. (**C**) Suggested binding mode of BEA in Pdr5, overlaid with the binding mode of R6G (white sticks) from the cryo-EM structure. Here, BEA places one of its benzyl moieties at a similar position as one of the benzene rings of R6Gs xanthene-like moiety. (**D**) Difference in the R6G binding site between the R6G and ADP-orthovanadate (AOV) bound Pdr5 structure (blue and green, respectively) viewed from above. The L572 and L573 containing helix in the AOV-bound structure moved towards the center of the R6G binding site and, thus, clashes with the suggested BEA binding mode. As a result, no binding mode of BEA was identified int the R6G binding pocket of the AOV-bound structure during the docking.

### In vitro measurement of substrate dependent H^+^ co-transport

In order to verify the results of the electrophysiological single molecule experiments, we investigated the time-dependent change of the bulk pH value in suspensions with Pdr5 enriched plasma membrane vesicles under conditions of active Pdr5 substrate transport with a bulk fluorescence assay. Since many fluorophores are substrates of Pdr5WT, we used a pH sensitive fluorescein derivate (see SI for further details) conjugated to dextran with a molecular weight of about 40 kDa thus excluding that the dye might be a substrate of Pdr_WT_. (17).

Furthermore, we visualized Pdr5 enriched plasma membrane vesicles using negative stain electron microsopy. Both yeast plasma membrane preparations of Pdr5_WT_ and Pdr5_EQ_ form typical uniform vesicular structures with an average diameter of 300-500 nm (Fig. 9A).

**Figure 9.**
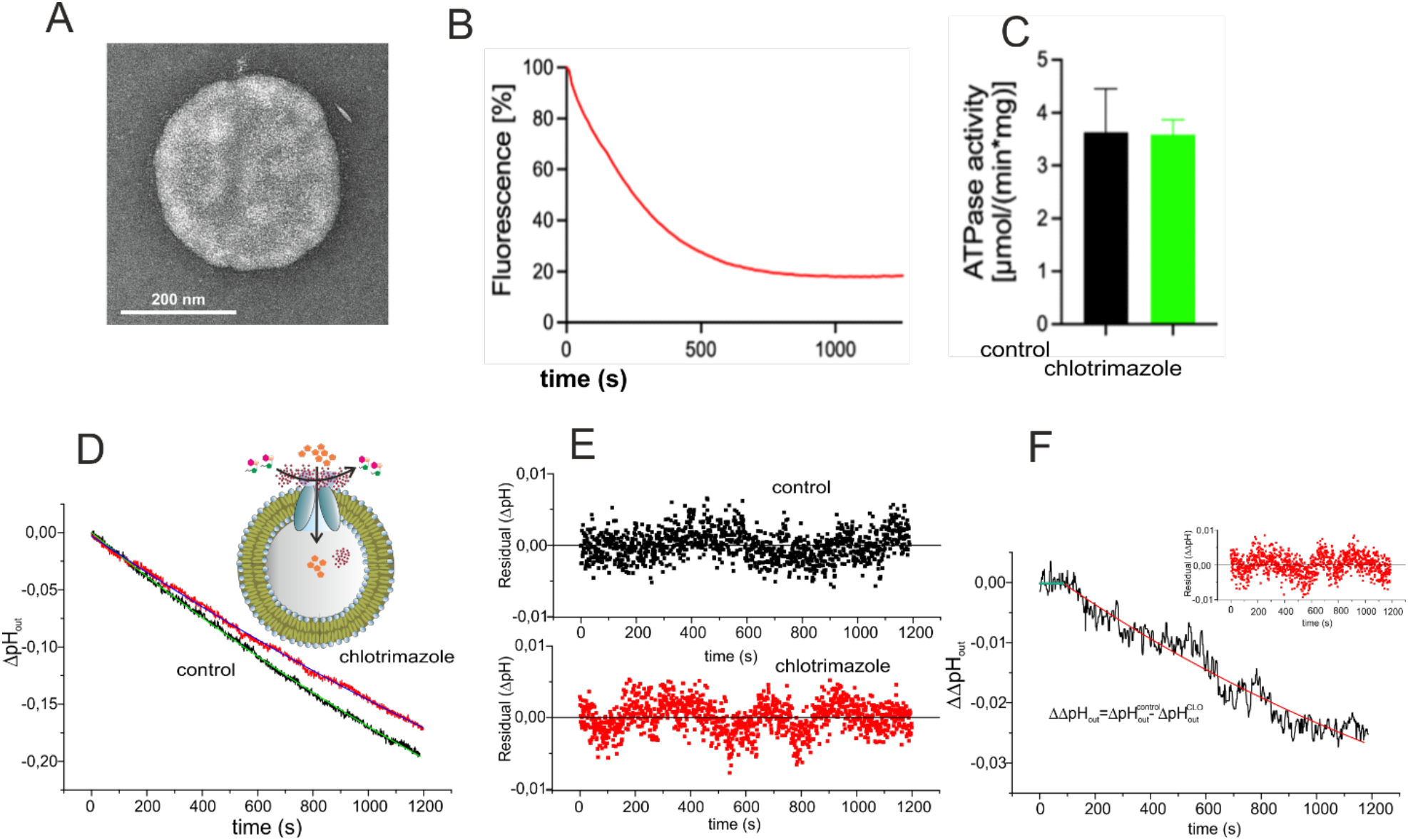
Fluorescence pH_out_ assay of ATP driven substrate transport in Pdr5_WT_ in yeast plasma-membrane-vesicles. (**A**) A representative negative stain micrograph of yeast membrane-vesicles enriched with Pdr5_WT_. Scale bar: 200 nm. **(B)** Control measument of R6G tranport in Pdr5_WT_ yeast plasma-membrane-vesicles under identical experimental conditions of the pH_out_ measurements, i.e. without additional external buffer. (**C**) Monitor of the Pdr5-specific ATPase activity of the Pdr5_WT_ membrane-vesicles under pH_out_ assy conditions in the absence (black bar) or in the presence of 1 μM clotrimazole (green bar). Data represents the average of six independent experiments with the error reporting the standrad deviation. p-value determined by an unpaired t-test implemented in Prism 9 (GraphPad Inc.) between both data sets is 0.859 indicating no significant differences in the respective ATPase activity. (**D**) Time dependent pH_out_ changes monitored by dextran(40kD)-fluorescein_out_ in the absence (control, black) or presence of 1 μM clotrimazole_out_ (red). Dextran(40kD)-fluorescein fluorescence was calibrated in the assay buffer containing additional KP_i_-buffer between pH6.5 and 8.0. For ease of visulaization, the time dependent change of dextran(40kD)-fluorescein_out_ fluorescence during the ATP induced transport was -with the help of the calibration curve -converted directly into ΔpH_out_ values and after averaging plotted as a function of time. Data represents the average of three independent experiments with the error reporting the standrad deviation. The solid lines (control, green; black, clotrimazole) represent the single exponential decay fit of the respective experimental ΔpH_out_ decay curves. **(E)** Residuals of the respective data-fit in (D). **(F)**Calculated difference 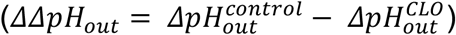 of the time-dependent ΔpH_out_ changes. Solid line shows the single exponential fit of the respective ΔΔ*pH_out_* curve and the inset the residuals of the fit.

These figures show that the stable yeast plasma membrane vesicle preparations meet the requirements to enable the investigation of bulk pH-changes mediated by transporter induced vectorial flux of substrates between the bulk phase and the inner lumenial space of the vesicles.

Second, we analyzed whether the low buffer capacity conditions required for the pH-assay might effect or even abolish Pdr5_WT_ mediated R6G transport in plasma membrane vesicles. As shown in Figure 9C, quenching of R6G fluorescence as read-out for Pdr5 transport activity was quantitatively not impaired and the ΔF_i_-equilibrium value reached approximately 20% residual fluorescence-intensity, which is well in the range of published values (17, 18). For ΔpH_out_ measurements, CLO was selected as substrate as it is at the given experimental conditions (in contrast to R6G) non-fluorescent and showed a V_rev_ = −57 ± 6.7 mV in the single channel recordings (Figure 6). Third, we checked wether the Pdr5-specific ATPase activity of the Pdr5 enriched membrane-vesicles under pH_out_ assay conditions were impaired in the presence of 1 μM clotrimazole outside. As shown in Figure 9D within error limits no differences between the ATPase activity of the control (black) and the one in the presence of 1 μM clotrimazole outside could be detected.

Furthermore we have recorded the time-dependent change of the bulk pH value (ΔpH_out_) in the suspension with Pdr5 enriched plasma membrane vesicles under conditions of active Pdr5 substrate transport. We observed a time dependent decrease of the dextran(40kD)-Fluorescein_out_ fluorescence intensity (ΔpH_out_). Three independent sample-recordings but with identical assay conditions were averaged and the observed time dependent fluorescence changes were with the aid of the calibration curve (see Materials and Methods) -converted directly into ΔpH_out_ values and plotted as a function of time. Figure 9E shows the timecourse of the ΔpH_out_ changes in the absence (black) and the presence of 1 μM clotrimazole (red). The solid lines in the respective recordings represent the results of the mono-exponential fit of the data and in Figure 9F the respective residuals of the fit are shown. As already obvious from the raw graph there is a significant difference in the time course of the decay curves. The fit of the single exponential ΔpH_out_ decay revealed time constants of *τ* = 2147.8 min^−1^ (control) and *τ* = 2699.7min^−1^ (CLO). Thus the relative ATP hydrolysis related H^+^-release per timeunit as shown in the decay curve of the control due to the uncoupled ATPase activity of Pdr5_WT_ (18), which releases one proton per hydrolyzed ATP, is reduced by a factor of 0.8 during CLO transport. Since however a CLO induced inhibition of ATPase activity has been ruled out experimentally (Figure 9F) the apparent lower amount of released H^+^ can in light of the electrophysiological results conclusively be explained by the substrate dependent co-transport of H^+^. The time course of the difference of ΔpH_out_ 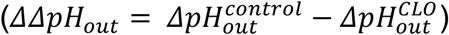 is shown in Figure 9G, the solid line represents the single exponential fit with a decay rate of *τ* = 1350 min^−1^ and the inset the respective residuals of the fit. Since we present strong evidence that this time dependent decrease of pH_out_ represents the rate of H^+^ cotransport into the vesicle lumen this results in an overall rate of *τ* ≅ 1350 min^−1^ for the H^+^-co-transport by Pdr5 in the vesicle suspension. In summary the results of the bulk fluorescence assay -which allowed us to determine – bulk pH changes in suspensions with Pdr5 enriched plasma membrane vesicles under conditions of active Pdr5 substrate transport-agree very well with the electrophysiological single molecule results and show that Pdr5 is an H^+^-cotransporter.

## Discussion

MDR ABC transporters have a broad range of substrates that are highly diverse in terms of structure and physico-chemical properties. There has been a long debate on how these membrane proteins confer resistance, i.e. how transporters reduce the drug concentration within a cell. Currently two models are favored: (i) the ‘drug pump model’, in which ABC transporters actively expel the compounds out of the cell is supported by numerous studies (36, 37) and (ii) the ‘altered partitioning model’, in which MDR ABC transporters confer resistance by alteration of the membrane environment through ion transport that ultimately results in lower drug uptake by the cell (7, 8, 12, 38). Essentially, the question arises whether all of the known substrates of MDR transporters are actually real substrates in terms of being actively transported out of the cell against the concentration gradient, or if these proteins mediate a decreased influx of the drugs by altering membrane properties, or whether both mechanisms are actually used.

With regard to these open questions, we tried to establish an *in vitro* transport assay for the well-studied yeast MDR ABC transporter Pdr5 (for a recent review see (16)). This transporter has been extensively studied in the last three decades and a robust *in vivo* transport assay based on the Pdr5- and ATP/Mg^2+^-dependent R6G quenching in plasma membrane preparation is available (17, 18). Despite all our efforts, we could not transfer the plasma membrane-based transport assay to proteoliposomes. Nevertheless, we also detected that the R6G transport in plasma membrane preparation was strongly dependent on artificial ion gradients (Figure 1), which is supported by other studies (25–27). Therefore, we turned to an electrophysiological *in vitro* approach, which enabled us to measure directly Pdr5-mediated ion conductivities under varying conditions. Previously, studies performed on MsbA and LmrA demonstrated that ion gradients benefit or are even indispensable for the activity of the respective ABC transporter, while ATP, the energy source of this protein family, seems to play only a regulatory role (9–13, 39). Moreover, in the case of LmrA, it remains elusive whether this pump works as an importer or exporter. Besides these complex, slightly contradicting studies, there are to our knowledge so far, no detailed electrophysiological analysis of MDR ABC transporters *in vitro*. On basis of the functionally purified, reconstituted Pdr5, and a well-studied ATPase deficient mutant as a control as well as known substrates of Pdr5 (R6G, KA, CLO and CHX (15, 17, 18, 20, 40)) and the potent Pdr5-specific inhibitor BEA, we were able to perform an extensive electrophysiological *in vitro* characterization of this MDR efflux pump.

In particular, we were able to demonstrate that Pdr5_WT_, but not the ATPase-deficient Pdr5_EQ_ mutant, generated a membrane potential that can only be the result of a substrate and Mg^2+^-ATP driven proton co-transport across the membrane. These data were also supported by *in vitro* measurements of bulk pHout changes with Pdr5 enriched plasma membrane vesicle suspensions. (Figure 9). Here, we observed an apparent significant slower ATP hydrolysis driven decrease in ΔpH_out_ if the substrate, CLO, was present. Considering that this time dependent decrease in ΔΔpH_out_ represents the rate of H^+^ cotransport into the vesicle lumen this results in an overall rate of *τ* = 22 s^−1^ for the H^+^-co-transport mediated by Pdr5 within the suspension of yeast plasma membrane vesicles.

This difference (ΔΔpH_out_) cannot be assigned to CLO induced alterations of the ATPase activity of Pdr_WT_ as, the Pdr5–ATPase activity remained the same within statistical error in the absence or presence of 1 μM CLO (Figure 9D). In contrast, Golin *et al*. reported a slight stimulation of the ATPase activity of Pdr5_WT_ in the presence of 1 μM CLO (41). Provided this would have been the case in our experiments, a stimulated ATPase activity in the presence of the substrate would result in higher rates of released H^+^ due to the stimulated hydrolysis of ATP. This would result in an even larger difference of the ΔΔpH_out_ decay-rate constant (Figure 9G). In summary our data demonstrate that bulk pHout changes with Pdr5 enriched plasma membrane vesicle suspensions induced by the ATP-hydrolysis drop significantly in the presence of substrate. We therefore conclude that Pdr5 is an ATP-dependent drug/H^+^ co-transporter that actively translocates substrates and generates an additional PMF through vectorial proton efflux. Our electrophysiological measurements allowed not only to prove that Pdr5 conducts ions and thereby follows the ‘altered partitioning model’, but also to demonstrate that it actively transports its substrates across the bilayer. Interestingly, the H^+^ co-transport occurs independently of the charge of the substrate and both, the cationic R6G as well as the neutral KA and CLO are transported in a drug/H^+^ co-transport fashion. The zero-current potentials for KA, CLO and R6G (V_rev_ = – 64 ± 8 mV, −57.3 ± 6.7 mV and – 47.2 ± 5.3 mV, respectively) clearly show that R6G is transported from cis to trans since due to the presence of R6G only on the cis-side the putative V_rev_ of the R6G^+^-ions contributes to the overall V_rev_, reducing it in comparison to the neutral KA and CLO. Additionally, using cationic and neutral substrates, we demonstrate that the measured H^+^ co-transport cannot be an artifact of the efflux of a protonated or cationic substrate and instead is a real H^+^ co-transport by Pdr5 in a Mg^2+^-ATP- and substrate-dependent manner. The recently described single particle cryo-EM structures of Pdr5 (34) do not yet led to a direct identification of amino acid residues potentially involved in protonation/deprotonation reactions leading to the establishment of a vectorial H^+^ translocation and finally the generation of an electrochemical gradient 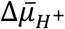. This situation is somehow reminiscent to LacY (42–44) and further experiments are required to pin-point the molecular pathway of proton transport.

The results presented in this study are also in line with the widely used R6G transport assay that was established for Pdr5 located in inside-out oriented plasma membrane vesicles (17, 18, 45, 46). R6G is a mitochondrial marker and a fluorescent membrane potential probe (27). The observed fluorescence intensity decrease in plasma membrane vesicles reflects the R6G and proton co-transport and thereby the membrane potential change during transport.

Another tested substrate, CHX, however, did not induce any proton gradient or change in conductance. In contrast to the majority of known Pdr5 substrates, CHX does not affect the ATPase activity of Pdr5 and is hydrophilic (18, 20). One can speculate that either a different transport route for this hydrophilic substrate exists, or that CHX is not a substrate at all. The observed resistance conferred by Pdr5 could be due to intracellular alkalization caused by Pdr5-mediated proton pumping as CHX undergoes degradation at basic pH (47).

An interesting feature of Pdr5 is that it can be voltage activated thus allowing high current fluxes across the membrane even in the absence of Mg^2+^-ATP or substrates. This activation occurred if Pdr5 was exposed to membrane voltages of V_h_ > +100 mV. In a physiological context with a yeast plasma membrane potential of about −82 mV (48) and an additional Pdr5 generated Δφ of about Δφ = −50 to −65 mV (V_rev, R6G_ = −47.2 ± 5.3 mV, V_rev, CLO_ = −57.3 ± 6.7 mV, and V_rev, KA_ = − 64 ± 8 mV), Pdr5 possibly activates itself into this high conductance state.

Next to the self-activation, the proton pumping properties of Pdr5 with multiple active copies would lead to the acidification in the trans compartment of our setup which reflects an acidic external pH in an *in vivo* setting. Moreover, drug/proton co-transport would temporarily significantly increase the membrane potential of the plasma membrane thus effecting the ion and solute homeostasis and the membrane morphology of yeast cells (49–51). This compares well with results that were obtained previously for the Pdr5 homologue Cdr1 and MDR1 *in vivo* as it has been speculated that MDR1 as well as Cdr1 may transport protons from the intra- to the extracellular space and thus alkalize the cytoplasm (8, 38). However, the complexity of the cell environment these *in vivo* experiments did not allow to unambiguously pinpoint observations to a single protein level nor for the experimenter to control all relevant critical cellular parameters. For Cdr1 it was hypothesized that the ABC transporter creates an additional PMF when it was expressed in *S. cerevisiae*. In this context it is important to note that in our experiments, the proton gradient does only occur in a strictly Mg^2+^-ATP and substrate dependent manner. This means that in the absence of a substrate, Pdr5 does not participate in pH homeostasis of the membrane even if ATPase activity is present. Therefore, the active proton pumping appears to be a second mode of action of this MDR ABC transporter.

Furthermore, we demonstrate that Pdr5 conducts K^+^ and Cl^−^ ions with a slight cation selectivity. This is not surprising as all known charged substrates of Pdr5, like for most other MDR ABC transporters, are cationic (15, 17, 37). We were also able to show that the membrane part of the nucleotide-free Pdr5 switches between at least three major confirmations as apparent from the three distinct subconductance states of which the main state is compatible with a pore diameter of 6.4 Å (52), which is within the range reported for other ABC transporters (53). Moreover, the broad distribution of open channel states indicates that Pdr5 has a rather flexible membrane part responsible for the translocation of its substrates. Additional experiments with the Pdr5 inhibitor BEA demonstrated that inhibition of the ATPase activity abolishes H^+^-co-transport and locks Pdr5 in an open conformation as indicated by the measured high conductance. Molecular docking suggests that this is mediated via BEA binding to the R6G binding site of Pdr5.

In conclusion, the presented data demonstrates that Pdr5 is an ATP-driven drug/proton co-transporter that, unlike LmrA or MsbA, does not use the PMF but creates a proton gradient across the membrane that results in self-activation and presents a novel mechanism of drug resistance.

## Materials and Methods

### Pdr5 Expression and Purification

Pdr5 was expressed and purified as described (20). Briefly, *S. cerevisiae* YRE1001 (18) cells were grown at 30 °C to a final OD_600_ of 3.5. Cells of 4 L YPD media were harvested and disrupted using glass beads. Total membranes were collected adjusted to 10 mg/ml total protein concentration with buffer A (50 mM Tris-HCl, pH 7.8, 50 mM NaCl, 10% (w/v) glycerol and solubilized for 1 h with 1% (w/v) trans-PCC-α-M under gentle stirring at 4 °C. Immobilized metal ion affinity chromatography of the His-tagged Pdr5 was performed with a Zn^2+^ loaded 1 ml HiTrap Chelating column (GE Healthcare) and elution was carried out with a step gradient using low and high histidine buffers (50 mM Tris-HCl pH 7.8, 500 mM NaCl, 10 % (w/v) glycerol, 0.003% (w/v) trans-PCC-α-M and 2.5 mM or 100 mM L-histidine. The elution fractions were collected, pooled and concentrated in a ultrafiltration unit (100 kDa MWCO). Size exclusion chromatography was performed on a Superdex 200 10/300 GL column (GE Healthcare) with buffer A containing 0.003 % (w/v) trans-PCC-α-M. All chromatography steps were performed on Äkta protein purification systems (GE Healthcare).

### Preparation of Pdr5 containing plasma membranes

Plasma membranes highly enriched in Pdr5_WT_ were prepared as described in (18).

### Negative stain electron microscopy

4 μl sample droplets (plasma membranes sample enriched in Pdr5WT or Pdr5EQ) were applied on freshly glow-discharged copper grids (Agar scientific; G2400C) covered by a thin, continuous carbon film. The sample was left for 2min on the grid before blotting, using filter paper (Whatman no. 4), subsequently washed with distilled water and stained with 10 μl 0.75% uranyl formate. Images were taken with a Tecnai Spirit electron microscope equipped with a LaB6 cathode operated at 120 kV. Digital electron micrographs were recorded with a 4k x 4k CMOS camera F416 (TVIPS) at a magnification of 67,000 x. (pixel size of 1.6 Å) using minimal dose conditions.

### Transport assays using Pdr5 containing plasma membranes

Transport assays were performed according to the protocol developed by Kolaczkowski and co-workers (17), using a FluoroLog III fluorescence spectrometer (Horiba). Isolated plasma membranes (30 μg of total protein content) were resuspended in 1 ml of transport buffer (50mM Hepes (pH 7.0), 10mM MgCl_2_, 150 nM R6G, and 10mM NaN_3_) and incubated at 30 °C. R6G Transport was initiated by addition of 5 mM ATP unless otherwise stated. The excitation wavelength was 529 nm (slit 1 nm) and the emission wavelength was 553 nm (slit 2 nm). For R6G transport data shown in Figure 9, no Hepes buffer was added to the assay medium.

### Single Channel Recordings from Planar Lipid Bilayers and Data Analysis

Planar lipid bilayer measurements using the Compact bilayer platform (Ionovation GmbH) were performed as described in detail previously (54). In brief: if not explicitly stated otherwise, symmetric conditions (250 mM KCl, 10 mM HEPES, pH 7.0) were used in cis and trans compartments. The denomination cis and trans corresponds to the half-chambers of the bilayer unit. Reported membrane potentials are always referred to the trans compartment. Bilayer fabrication was performed on PFTE film at a 100 μm prepainted (1 % hexadecane in n-hexane) aperture with a Phosphatidylcholine (18:1) (PC) / Phosphatidylethanolamine (18:1) (PE) (7:3 ratio) lipid mixture (both lipids were purchased from Avanti Polar Lipids) in n-pentane using the “thinning method” (54). Stock solutions of purified Pdr5 with typically 0.5-2.5 mg/ml contained in a buffer of 50 mM Tris-HCl pH 7.8, 50 mM NaCl, and 0.003% (w/v) trans-PCC-α-M were added to the cis compartment under slight stirring to a final protein concentration of ~1μg/ml to 2μg/ml.

Ion channel currents were recorded using an EPC 10 USB amplifier (HEKA Elektronik GmbH) in combination with the Patchmaster data acquisition software (HEKA Elektronik GmbH). For data acquisition a sampling rate of 5kHz (voltage ramps) and 10 kHz (continuous recording) was used and the data were further analyzed using the Origin package (Origin Lab) and the MATLAB (MathWorks) based Ion-channel-Master software developed in our laboratory (54).

### Molecular Docking

For the molecular docking BEA was drawn and converted into 3D using the ChemDraw19 suite. BEA was subsequently docked into four different Pdr5 cryo-EM structures utilizing a combination of AutoDock as a docking engine and the DrugScore2018 distance-dependent pair-potentials as an objective function. For this, the co-resolved ligands were removed and the structures aligned on the R6G bound Pdr5 structure using Pymol 2.4 (55). The DrugScore scoring grids were calculated for each Pdr5 structure twice: It was centered on BEA, which was manually docked into either the R6G binding site or the space between the TMDs and NBDs of Pdr5. Both these initial BEA structures were docked into either grid to minimized starting structure bias. During docking, default parameters were used, except for the clustering RMSD cutoff, which was set to 2.0 Å(56–58). Binding modes were considered valid if they were contained in the largest cluster with the most favorable docking energies, which comprised at least 20 % of all docking poses.

### ATPase measurements in Pdr5-enriched plasma membranes

Plasma membranes were prepared, and ATPase activity was determined as described previously (18). Briefly, oligomycin (OM) sensitive ATPase activity of plasma membrane fractions was measured by a colorimetric assay. Isolated plasma membranes (0.2 μg per well) were incubated with 5 mM ATP, 5 mM MgCl_2_ in 300 mM Tris-glycine buffer (pH = 9.0) in a final volume of 100 μL. Pdr5-specific ATPase activity was measured in presence or absence of 1 μM clotrimazole. To reduce background, 0.2 mM ammonium molybdate, 50 mM KNO_3_, and 10 mM NaN_3_ were added. In a control reaction, OM (20 μg mL^−1^) was added to the assay to determine the OM-sensitive ATPase activity. After incubation at 30 °C for 20 min, the reaction was stopped by adding 25 μL of the sample to 175 μL of ice-cold 40 mM H_2_SO4. The amount of released inorganic phosphate was determined by a colorimetric assay, using Na_2_HPO_4_ as standard.

### Fluorescence pH measurements

pH measurements were performed accordingly to the R6G transport in Pdr5-enriched plasma membranes, i.e. R6G was substituted by 2 μg/ml Fluorescein conjugated Dextran (40,000 MW, see SI for further details). The excitation wavelength was 494 nm (slit width 1 nm) and the emission wavelength was 521 nm (slit width 2 nm). The pH measurements were performed in absence or presence of 1 μM clotrimazole. For data evaluation a calibration curve with a R-squared value of 0.94 was generated using 50 mM KP_i_ buffer with defined pH values between 6.5 and 8.0.

## Acknowledgments

We thank Prof. Oliver Ebenhöh (Heinrich Heine University Düsseldorf), Prof. Alexej Kedrov and all members of the Institute of Biochemistry for stimulating discussions. We also thank Prof. Mathias Winterhalter (Jacobs University Bremen) for support. A.S.-H. was funded through ERC Advanced grant VisTrans (ID: 742210). The Center for Structural Studies is funded by the Deutsche Forschungsgemeinschaft (DFG Grant number 417919780 to S.S.).

## Acknowledgments

The authors declare no financial and non-financial competing interest.

## Notes

### Competing Interest Statement

The authors have declared no competing interest.

### Summary of Updates

Added new experimental data (Figure 9)

