## Supplementary data for "A new twist in ABC transporter mediated multidrug resistance – Pdr5 is a drug/proton co-transporter"

**SI Appendix**

**Table of Contents**

**Figure S1:** All point amplitude histograms for single channel recordings from a $Pdr5_{WT}$ containing bilayer shown in Figure 2C.
**Figure S2:** Ion selectivity of the activated $Pdr5_{WT}$ channel.
**Figure S3:** Comparison of experimental and calculated fits for the reversal potentials.
**Table S1**
**Table S2**
**Table S3**
**Table S4**
**Methods**
**References**

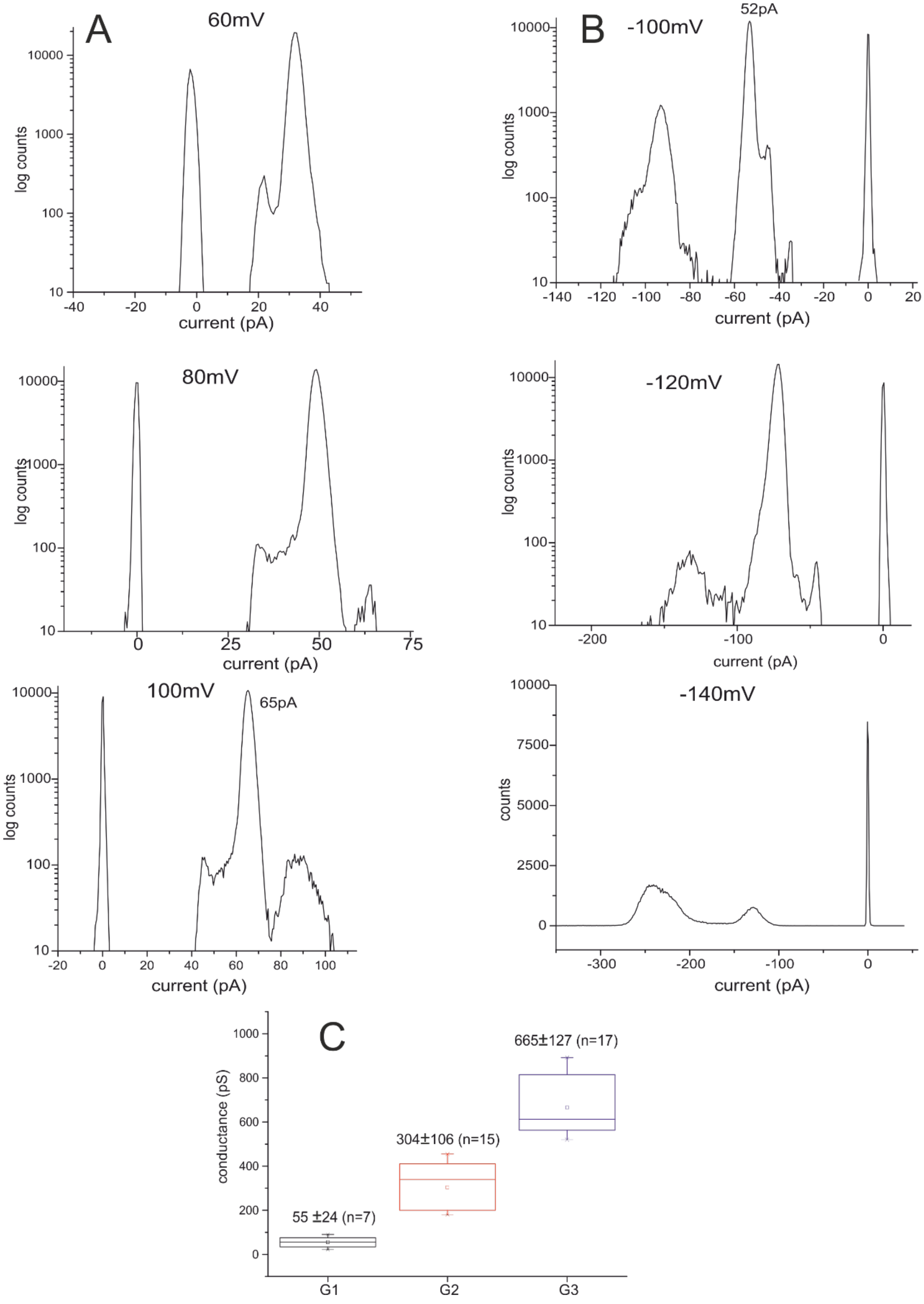


**Figure S1. All point amplitude histograms for single channel recordings from a $Pdr5_{WT}$ containing bilayer shown in Figure 2C.** (**A**) Histograms from recordings at the indicated positive $V_h$ (**B**) Histograms from recordings at the indicated negative $V_h$. **(C)** Box-chart plot for the three mean $Pdr5_{WT}$ conductance states from N=5 bilayer measurements derived from the mean variance analysis of n-recordings.

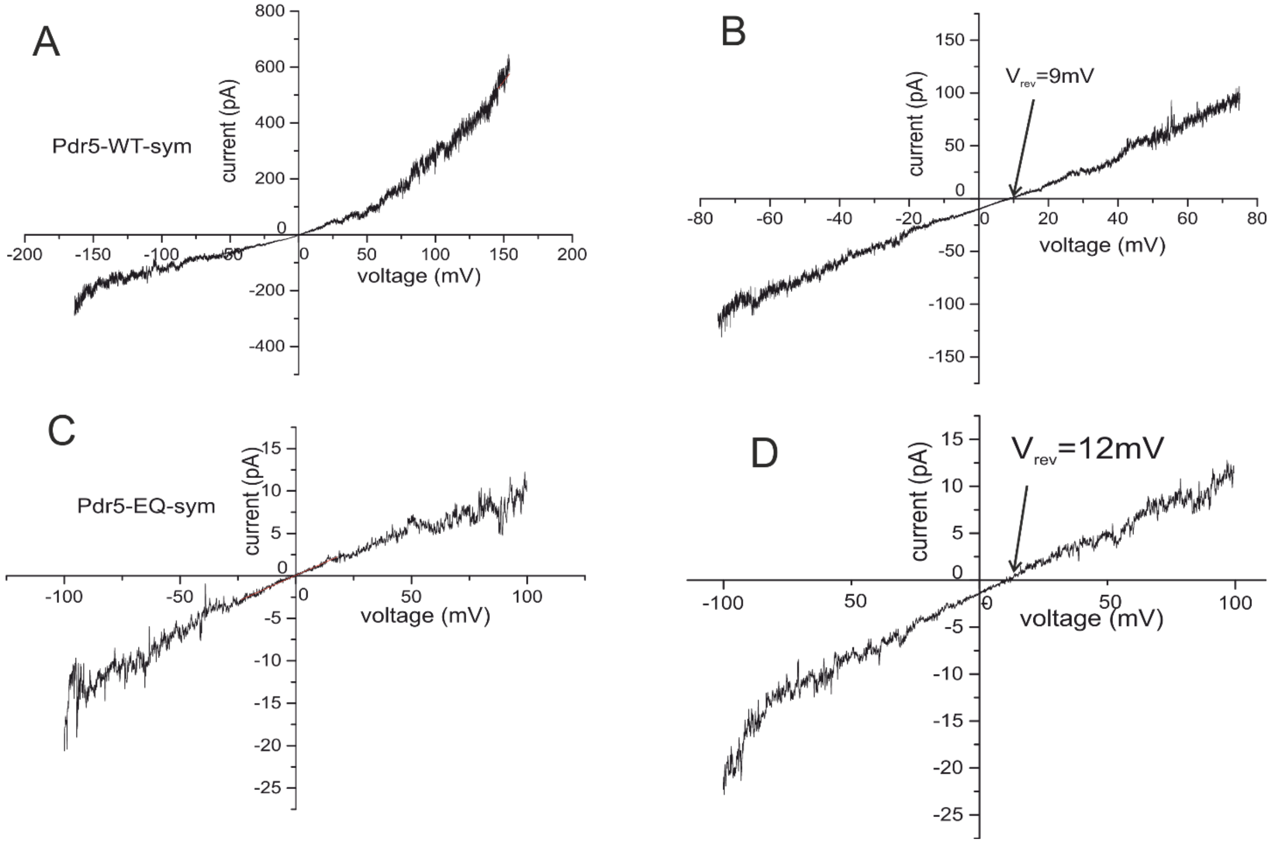


**Figure S2 Ion selectivity of the activated Pdr5$_{WT}$ and Pdr5$_{EQ}$ channel.** (**A**) Current recording from a bilayer containing multiple active copies of the reconstituted voltage activated Pdr5$_{WT}$ in response to a voltage ramp from -150 mV to +150 mV in symmetrical cis/trans (250 mM KCl, 10 mM HEPES, pH 7.0) buffer. Pdr5$_{WT}$ always displayed a rectifying current voltage relation with significant higher currents at positive $V_h$. (**B**) Asymmetric buffer conditions (590 mM/250 mM KCl (cis/trans)) led to $V_{rev}$ of +9 mV.
(**C**) Current recording from a bilayer containing multiple active copies of the reconstituted voltage activated Pdr5$_{EQ}$ in response to a voltage ramp from -150 mV to +150 mV in symmetrical cis/trans (250 mM KCl, 10 mM HEPES, pH 7.0) buffer. Pdr5$_{WT}$ always displayed a rectifying current voltage relation with significant higher currents at positive $V_h$.
(**D**) Asymmetric buffer conditions (590 mM/250 mM KCl (cis/trans)) led to $V_{rev}$ of +12 mV.

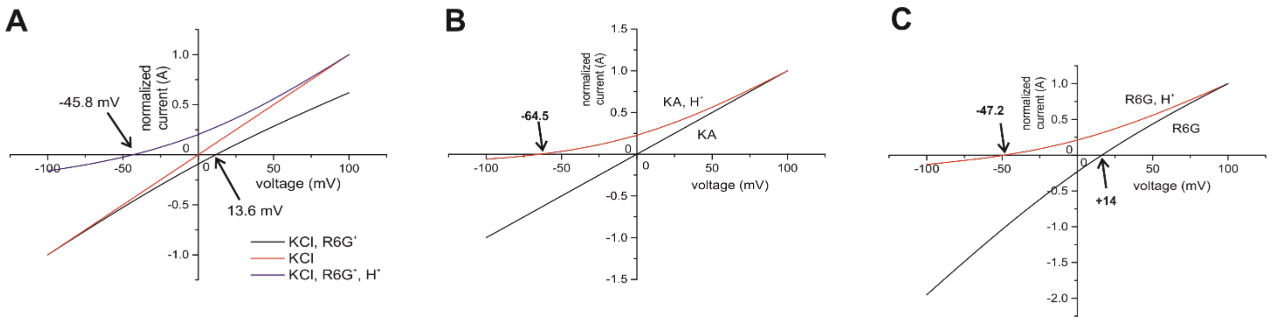


**Figure S3 Comparison of experimental $V_{rev}$ values with calculated current voltage relations.** (**A**) Current-voltage relations calculated by the GHK current equations using experimental ionic conditions and the respective permeability ($P_x$) as variable (see below for details) for 3 cases. **Red:** KCl (only permeation of $K^+$ and $Cl^-$ ions), **Black:** KCl-R6G (permeation of $K^+$ and $Cl^-$ ions and active transport of $R6G^+$ ions), **Blue:** KCl-R6G-$H^+$ (permeation of $K^+$ and $Cl^-$ ions and active co-transport of $R6G^+$ and $H^+$ ions). Currents were normalized to $i_{max}$. For more details see Methods section and SI. (**B**) Fit of the experimental reversal potentials (**red**) and comparison to the calculated fit of the reversal potentials if no protons are co-transported **(black)** for KA. (**C**) Fit of the experimental reversal potentials **(red)** and comparison to the calculated fit of the reversal potentials if no protons are co-transported **(black)** for R6G.

**Table S1.** MS analysis of purified Pdr5.

| Accession | | Score | Coverage | # Proteins | # Unique Peptides | # Peptides | # PSMs | # AAs | MW [kDa] |
|---|---|---|---|---|---|---|---|---|---|
| P33302 | PDR5 | 146072,5 | 58,77 | 1 | 68 | 82 | 652 | 1511 | 170,3[illegible] |
| Q04182* | PDR15* | 28336,6 | 10,2 | 1 | 2 | 16 | 133 | 1529 | 172,1[illegible] |
| Q08970 | MMT2 | 2633,2 | 20,45 | 1 | 8 | 8 | 13 | 484 | 52,4[illegible] |
| Q03218 | MMT1 | 1233,8 | 11,37 | 1 | 5 | 5 | 6 | 510 | 56,1[illegible] |
| P38248 | ECM33 | 962,3 | 10,96 | 1 | 4 | 4 | 5 | 429 | 43,7[illegible] |
| P60010 | ACT | 885,9 | 10,4 | 1 | 4 | 4 | 5 | 375 | 41,6[illegible] |
| P12904 | AAKG | 796,5 | 13,98 | 1 | 4 | 4 | 5 | 322 | 36,3[illegible] |
| P05030 | PMA1 | 794,1 | 5,23 | 2 | 4 | 4 | 4 | 918 | 99,5[illegible] |
| P22146 | GAS1 | 742,4 | 10,91 | 1 | 5 | 5 | 5 | 559 | 59,5[illegible] |
| Q08954 | YP199 | 734,5 | 15,83 | 1 | 4 | 4 | 5 | 240 | 26,7[illegible] |
| P0CG63 | UBI4P | 714,6 | 52,49 | 3 | 4 | 4 | 5 | 381 | 42,8[illegible] |
| P20107 | ZRC1 | 647,4 | 6,33 | 1 | 2 | 2 | 3 | 442 | 48,3[illegible] |
| P46992 | YJR1 | 613,8 | 13,38 | 1 | 4 | 4 | 4 | 396 | 42,9[illegible] |
| P25087 | ERG6 | 576,9 | 12,27 | 1 | 3 | 3 | 3 | 383 | 43,4[illegible] |
| P32798 | COT1 | 550,1 | 7,97 | 1 | 3 | 3 | 3 | 439 | 48,1[illegible] |
| P51996 | YPT32 | 405,1 | 14,41 | 1 | 3 | 3 | 3 | 222 | 24,5[illegible] |
| P32476 | ERG1 | 326,8 | 3,83 | 1 | 2 | 2 | 2 | 496 | 55,0[illegible] |
| P06782 | SNF1 | 325,5 | 2,53 | 1 | 2 | 2 | 2 | 633 | 72,0[illegible] |
| P22202 | HSP74 | 311,2 | 4,52 | 2 | 2 | 2 | 2 | 642 | 69,6[illegible] |
| Q08193 | GAS5 | 297,9 | 5,79 | 1 | 2 | 2 | 2 | 484 | 51,8[illegible] |
| P02994 | EF1A | 276,4 | 4,15 | 1 | 2 | 2 | 2 | 458 | 50,0[illegible] |
| P29704 | FDFT | 271,1 | 3,6 | 1 | 2 | 2 | 2 | 444 | 51,6[illegible] |
| P32623 | CRH2 | 268,0 | 4,28 | 1 | 2 | 2 | 2 | 467 | 49,8[illegible] |

*peptides that were identified under PDR15 are also found in PDR5

**Table S2.** Conductance parameter of non-activated $Pdr5_{WT}$ (Figure 5D-F) .

| | **$Pdr5_{WT}$** | **$Pdr5_{WT}$ -ATP** | **$Pdr5_{WT}$ -ATP-R6G** |
|---|---|---|---|
| **$G_{sl}$ (pS) (+$V_h$)** | 0.72 | 1.8 | 6.7 (19.2) |
| **$G_{sl}$ (pS) (-$V_h$)** | 19.2 | 21,4 | 31 |
| **$V_{rev}$ (mV)** | 0 | 0 | -58 |
| **Rectification** | 26.7 | 11.9 | 4.6 (1.6) |

**Table S3.** BEA induced conductivity across an empty bilayer (Figure 7A-C).

| | **Control** | **BEA (3.3μM)** | **BEA (9.9μM)** |
|---|---|---|---|
| **$G_{sl}$ (pS)** | 1.3 (Figure 4C) | 4.2 (Figure 4B) | 20.8 (Figure 4A) |
| **$V_{rev}$ (mV)** | 0 | 0 | 0 |

**Table S4.** Conductance parameters of $Pdr5_{WT}$ in the presence of $Mg^{2+}$-ATP, R6G and BEA (Figure 7D-G).

| | **Blank** | **Pdr5-$Mg^{2+}$-ATP** | **Pdr5-$Mg^{2+}$-ATP/R6G** | **Pdr5-MgATP/R6G-BEA (1.7μM)** |
|---|---|---|---|---|
| **$G_{sl}$ (pS)** | 1.2 | 2.4 (Figure 7E) | 7.7 (Figure 7F) | 51(Figure 7D) |
| **$V_{rev}$ (mV)** | 0 | 0 | -51 | 0 |

### Methods

After addition of Pdr5 to the cis compartment, channel insertion into the bilayer occurred unidirectional as deduced from the rectification characteristics of the current-voltage ramps. Gating transition amplitudes were analyzed and subsequently filtered for dwell times exceeding five times the sampling interval to exclude incompletely resolved gating transitions from the analysis (McManus, Blatz, & Magleby, 1987). When required and applicable, currents traces with amplitudes well below 10pA were filtered by a Savitzky-Golay filter (Gorry, 1990; Savitzky & Golay, 1964) to enhance the S/N ratio. Single channel analysis was performed essentially as described in detail (Harsman, Bartsch, Hemmis, Kruger, & Wagner, 2011). In brief, we used the robust mean variance approach (Patlak, 1988, 1993) that allows rigorous unbiased analysis of single channel recordings. For this, we had developed a Mathlab-based program that allows the automatic current-amplitude and channel state dwell time analysis of single channel recordings (Bartsch, Harsman, & Wagner, 2013).

### Determination of the ion selectivity and conductance sates of the voltage activated Pdr5 channel.

After voltage activation, reconstituted $Pdr5_{WT}$ always displayed a rectifying current voltage relation with significant higher currents at positive membrane voltages no matter whether single active cannels or multiple active channel currents were detected. Under asymmetric buffer conditions (590 mM/250 mM KCl (cis/trans)) a reversal potential of $V_{rev}$ = +9 mV ± 2.8 (n=3) was observed for the $Pdr5_{WT}$ , while for the $Pdr5_{EQ}$ mutant we obtained $V_{rev}$ = +13 mV ± 2.2 (n=3) with the same gradient (details not shown). Using the GHK approach the corresponding relative permeability for the $Pdr5_{WT}$ and the $Pdr5_{EQ}$ mutant are $P_K$+/ $P_{Cl}$-=2.5:1 and $P_K$+/ $P_{Cl}$-=3.5:1 respectively.

After voltage activation, reconstituted $Pdr5_{WT}$ displayed membrane potentials above ±100 mV and complex, fast channel gating (Figure 3). We analyzed the single channel recordings using the mean variance analysis, which allows an unbiased determination of ion-channel conductance states (Patlak, 1988, 1993). For this we applied the Matlab based Ion-Channel-Master program (Bartsch et al., 2013).

The $Pdr5_{WT}$ channel displayed fast flickering gating transitions from the closed to the open state with different open channel amplitudes (see expansion plot in the lower part of Figure 3C,D and Figure S1). From this results the question arises whether these different conductance states belong to a single $Pdr5_{WT}$ transporter or whether the particular channel open states reveal inhomogeneous open channel amplitudes of simultaneous channel opening or closing from different channels. The time-course of the gating events (see marked events) clearly indicates that the observed complex gating to different open states cannot be due to the presence of two or more active channels with simultaneous gating. The probability of to observe 2 simultaneous gating transitions within the sampling window of 100µs is: W= $(\tau_{s.})^2 = 1 \cdot 10^{-8} s$. Thus in 3 years and 21 days measuring time, the probability (W) of a simultaneous gating in this time window would be W = 1. The mean variance plot in Figure3E,F shows directly the gating transitions between 2 current amplitudes in the form of a square parabola.

We collected similar set off data $(\text{current recordings} \rightarrow histogram \rightarrow mean - variance)$ from N=5 bilayer with n ≥ 7 recordings with $V_h = \pm 60$ to $V_h = \pm 140$ mV. It turned out that the gating transitions could be grouped into three different Pdr5 conductance states: $G_1 \cong 55 \pm 24\, pS,\ G_2 \cong 304 \pm 106\, pS,\ G_3 \cong 665 \pm 127\, pS.$ (see box-chart-plot in Figure S1C).

**Determination of the substrate induced reversal potential shift with the reconstituted non activated Pdr5.**

After reconstitution into a planar bilayer, $Pdr5_{WT}$ induced small ion currents and we tested whether the substrates Ketoconazole (KA) and Rhodamine 6G (R6G) in the presence of $Mg^{2+}$-ATP induces shifts in a similar way as with the voltage activated $Pdr5_{WT}$.

In these measurement, addition of KA to the cis compartment induced a drastic shift of the reversal potential of $V_{rev}$ = -70mV. In four different bilayer experiments we obtained an average of $V_{rev}$ = -64 ± 8 mV (n=4).

Similar experiments were conducted with R6G as a substrate. Starting with the control bilayer (Figure 3D), we added $Pdr5_{WT}$ to the cis compartment of the same bilayer and observed spontaneous insertion of $Pdr5_{WT}$ into the bilayer, without applying prolonged high voltage, a small rectifying current (Figure 1D). After subsequent addition of $Mg^{2+}$-ATP to cis/trans (Figure 3E) and in the presence of 330nM R6G (Figure 3F) we observed again a significant shift in the reversal potential (Figure 3F). This shows that in the presence of R6G an additional cation flux from cis to trans with $\Delta V_{rev} = -58\,mV$ must have occurred (Figure 6F). In different bilayer experiments at symmetrical 250 mM KCl, 10 mM HEPES, pH 7.0 buffer with $Mg^{2+}$-ATP (cis) and R6G (cis) as a substrate we observed a shift of the reversal potential of $V_{rev} = -47.2 \pm 5.3 mV\ (n = 12)$.

**Background of the electrophysiological permeation assay (GHK approach).**

The Goldman-Hodgkin-Katz approach is by far the most commonly used framework to describe ion permeability and selectivity of membranes (Goldman, 1943; Hodgkin & Katz, 1949). Beside the principal difficulties underlying the macroscopic GHK constant field theory, which assumes independent movement of the ions through membrane pores (see references (Corry, Kuyucak, & Chung, 2000; Moy, Corry, Kuyucak, & Chung, 2000; Syganow & von Kitzing, 1999) for a detailed discussion) it has been demonstrated that the methodology can be used to obtain reliable semi-quantitative measures for permeation of charged drugs through membranes (Hille, 2001).

We were interested in obtaining information on the selectivity of the membrane transport mediated by the ABC transporter $Pdr5_{WT}$. For this, we used high-resolution electrophysiological recordings from planar bilayer containing the reconstituted $Pdr5_{WT}$. In practice, this method can be used to measure ion currents across the membrane up to a maximum resolution of 100 fA with a maximal bandwidth of 1 kHz (Hamill, Marty, Neher, Sakmann, & Sigworth, 1981). In the case of ion channels, for example, a channel with a conductance of G = 100pS the current through the open channel at $V_h$ = 100mV is i = 10pA corresponding to a turnover of $n = 6.2 \cdot 10^7 ions/s$. Therefore, by this method the current through a single ion channel molecule can be measured. The situation with $Pdr5_{WT}$ is different: from the ATP turnover of approx. 6 ATP / s (Ernst et al., 2008) which in the worst case is connected to the same turnover (stoichiometry 1:1) of a charged molecule across the membrane, we obtain a conductance of G = 10 fS and a current i = 0.001 fA ($V_m$=100mV). It is therefore obvious that even under the most optimal conditions, ion currents across the membrane mediated by individual ABC transporters cannot be resolved. On the contrary, in such a case of electrophysiological measurements, it is necessary to rely on the largest possible number of transporters being active.

A planar bilayer with a radius of 100µm has an area of approx. $F = 3 \cdot 10^{-4} cm^2$, which implies that the bilayer with 300 active $Pdr5_{WT}$ (see above) would have a specific conductivity of roughly $G_m = 10 \mu S/cm^2$. When applying a voltage of $V_m = +100mV$ this results in a total flux of about $n = 6.2 \cdot 10^7 ions/s$ to adjust the new membrane equilibrium. Moreover, from the specific resistance and specific capacity of the planar bilayer one can calculate that the new equilibrium states are realized with an exponential time constant of about 1ms. In other words*, at very small membrane currents the measurement of zero-current potentials largely reflects the relative current fluxes across the membrane rather than absolute concentrations in the cis and trans compartments* and is therefore an excellent method to study the properties of the ion fluxes mediated by transporters with low turnover rates across the membrane.

During the electrophysiological experiments with $Pdr5_{WT}$ reconstituted into planar bilayer, however, it turned out that $Pdr5_{WT}$ can be transferred by exposure to higher voltages ($V_m \geq \pm 100mV$) to function in an ion channel mode capable to conduct ions at high rates with a main conductance state of $G_2 \cong 300pS$ and subconductance states of $G_1 \cong 55pS$, and $G_3 \cong 665pS$. In addition, we observed that BEA binding to $Pdr5_{WT}$ opened in a concentration dependent manner activated $Pdr5_{WT}$ channels (Figure 7B,C) in a saturating fashion (details not shown) allowing an approximate estimate of the total number of $Pdr5_{WT}$ incorporated into the bilayer. On the other hand, we also detected very small membrane currents after incorporation of $Pdr5_{WT}$ into the planar bilayer without voltage activation (see Figure 3D-I ). These small currents revealed in asymmetric buffer (590 mM KCl (cis)/250 mM KCl (trans)) a reversal potential of $V_{rev}$=11mV (details not shown) a similar value as obtained for the high conductance form of $Pdr5_{WT}$ (see Figure S2B). This shows that also in this case the currents are likewise carried by $K^+$ and $Cl^-$ ions with almost the same permeability ratio of $\frac{P_{K^+}}{P_{Cl^-}} = 3.3$ as in the $Pdr5_{WT}$ high conductance states. One interesting estimate concerning the turnover for cations through $Pdr5_{WT}$ in the low turnover transporter mode can be deduced from measurements with the activated form (details not shown) and a BEA titration of bilayer yielding a maximal slope conductance of ~160 nS. Considering the mean conductance of $G_2$ = 300 pS, this means that the bilayer contained ~533 copies of the active $Pdr5_{WT}$. With the slope conductance of $G_{sl}(\mathrm{pS})$ (-$V_h$) = 19 pS (Table S1) we obtain a *"native"* conductance of $Pdr5_{WT}$ $G \cong 35\ fS$ in 250 mM KCl solution.

To characterize the ion fluxes mediated by the activated and non-activated $Pdr5_{WT}$ we employed the following experimental conditions for bilayer containing an unknown number for $Pdr5_{WT}$.

**Permeability $Pdr5_{WT}$:** $P_{K^+} = 2.5;\ P_{Cl^-} = 1;\ P_{R6G^+}; P_{KA}; P_{CLO} = variable;\ P_{H^+} = variable$
**Cation:** $z_{K^+} = 1;\ c_{K^+cis} = 250\ mM; c_{K^+\,trans} = 250\ mM$
**Anion:** $z_{Cl^-} = -1.0;\ c_{Cl^-cis} = 250mM;\ c_{Cl^-trans} = 250mM$
**Substrate KA:** $z_{KA} = 0;\ c_{AB^-cis} = 330nM;\ c_{KA-trans} = 00mM$
**Zero-current potential KA:** $V_{rev} = -64.5\ mV\ (experimental\ value\ n = 4))$
**Substrate CLO:** $z_{CLO} = 0;\ c_{CLO\ \ cis} = 1\ \mu M;\ c_{CLO-trans} = 00mM$
**Zero-current potential CLO:** $V_{rev} = -57\ mV\ (experimental\ value\ n = 4))$
**Protons:** $z_{H^+} = 1;\ c_{H^+cis} = 8 \cdot 10^{-9} M(pH\ 8.1)\ ; c_{H^+\,trans} = 1.04 \cdot 10^{-6} M(pH\ 5.98)$
**Substrate R6G:** $z_{R6G^+} = 1;\ c_{AB^-cis} = 330nM;\ c_{KA-trans} = 00mM$
**Protons:** $z_{H^+} = 1;\ c_{H^+cis} = 7.94 \cdot 10^{-9} M(pH\ 8.1)\ ; c_{H^+\,trans} = 9.3 \cdot 10^{-7} M(pH\ 6.03)$
**Zero-current potential R6G:** $V_{rev} = -47.2\ mV\ (experimental\ value\ (n = 12)$

Assuming electrical recording from a bilayer with 2mM $Mg^{2+}$-ATP (cis/trans) and the above concentrations in the cis and trans compartment and considering that the assumptions of the GHK-theory are valid under the applied conditions we can use equations (1 to 6 below) to calculate the expected current voltage relation for the above bilayer membrane containing an unknown number of active $Pdr5_{WT}$ channels (Figure S3).

**GHK-current equations**

1. $I_x(V, P_X, z, c_{cis}, c_{trans}) = P_x z^2 \frac{VF^2}{RT} \cdot \frac{\left(c_{x,cis} - c_{x,trans} exp\left(\frac{-zFV}{RT}\right)\right)}{1 - exp\left(\frac{-zFV}{RT}\right)}$
2. $I_{K^+}(V) = I(V, P_{K^+}, z_{k^+}, c_{K^+cis}, c_{K^+trans}),$
3. $I_{CL^-}(V) = I(V, P_{Cl^-}, z_{Cl^-}, c_{Cl^-cis}, c_{Cl^-trans})$
4. $I_{R6G^+}(V) = I(V, P_{R6G^+}, z_{R6G^+}, c_{R6G^+cis}, c_{R6G^+trans})$
5. $I_{H^+}(V) = I(V, P_{H^+}, z_{H^+}, c_{H^+cis}, c_{H^+trans})$
6. $\sum I(V) = I_{K^+}(V) + I_{Cl^-}(V) + I_{R6G^+}(V) + I_{H^+}(V)$

Using equation 1-6 we calculated for the ionic conditions given above the corresponding current-voltage relations using a Mathcad (PTC-Software) based routine. Starting with ketaconazole addition to the cis compartment in the presence of $Mg^{2+}$-ATP however we have to propose that the reconstituted $Pdr5_{WT}$ initiates active substrate and $Mg^{2+}$-ATP driven transport of $H^+$ ions from cis to the trans compartment ($H^+$ co-transport) to explain the observed negative $V_{rev}$ values (Figure 2F, Figure3B, C). Active transport means that the permeability (P) approaches in the GHK formalism towards infinity. Starting for KA at pH 7 (cis/trans) we obtained by generating a pH gradient through iteration the "new" equilibrium concentration of $H^+$ ions in the cis and trans compartment after initiation of transport with the correponding value of $P_{H^+}$ $(P = 10^6\ cm^3/s)$ and with the now fixed ΔpH we obtain the value for $P_{R6G^+}$ $(P = 2 \cdot 10^5\ cm^3/s)$. Since KA induced $H^+$-pumping by Pdr5 is expected to be in the same order of magnitude as for R6G it appeared justified to use the values $(P_{H^+,}\ \ c_{H^+cis} = 8 \cdot 10^{-9} M\ ; c_{H^+\,trans} = 1.04 \cdot 10^{-6} M$ obtained for KA to fit the observed reversal potential obtained with R6G as substrate. Provided $Mg^{2+}$-ATP driven R6G transport would occour without $H^+$ pumping at the given ionic conditions (cis/trans) we should have observed $V_{rev} \cong +14\ mV$ (Figure S4C) which was not the case. However our calculations show that $R6G^+$ and $H^+$ ions are co-transported at a relative permeability of $P_{H^+}/P_{R6G^+} \cong 5$.

### Fluorescence pH measurements

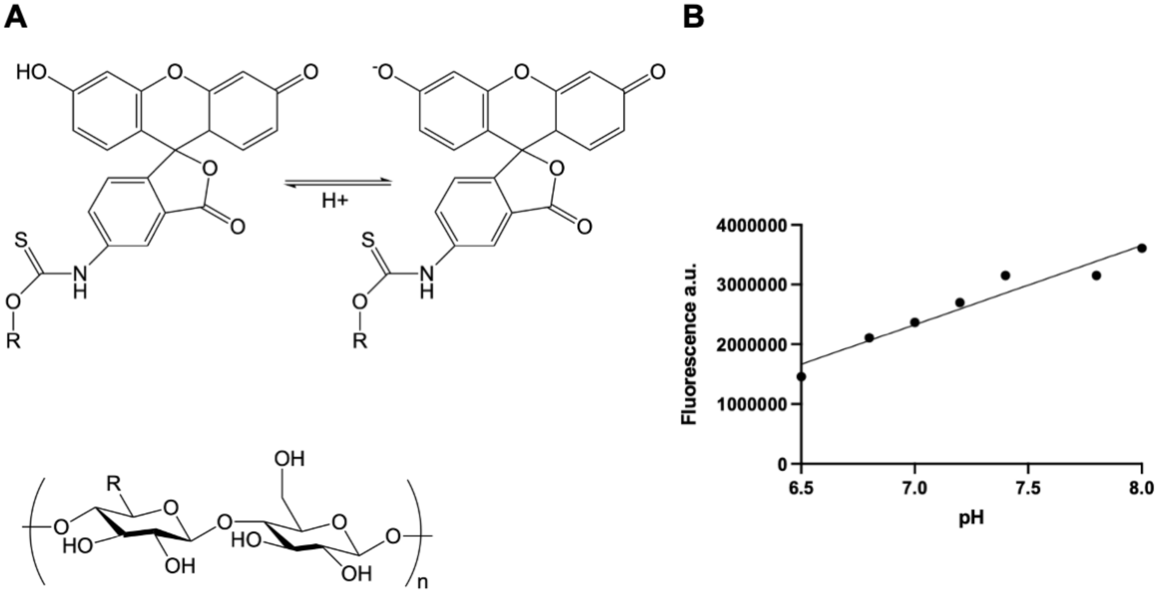


**Figure S4. Chemical structure and calibration curve of dextran(40kD)-fluorescein. (A)** Chemical structure of neutral and monoanionic fluorescein coupled to dextran**.** The building block of dextran is shown in the lower panel. The number of building blocks (n) accounts to a molecular weight of approximately 40 kDa. **(B)** Calibration curve of the pH dependent fluorescence-intensity-changes of dextran(40kD)-fluorescein in 50 mM $KP_i$. The fluorescence intensity was measured at an excitation wavelength of 494 nm (1 nm slit) and emission wavelength of 521 nm (2 nm) and plotted against the pH. The linear regression line was generated with the GraphPad Prism 9 software with the following equation: $y = 1324753x – 6945804$ ($R^2 = 0.94$).
